# Spectral Graph Features for Reference-free RNA 3D Quality Assessment

**DOI:** 10.64898/2026.04.06.716854

**Authors:** Ying Zhu, Huaiwen Zhang, Vince D. Calhoun, Yuda Bi

## Abstract

**Motivation:** Existing RNA 3D structure quality assessment (QA) methods rely on local geometric descriptors or statistical potentials that evaluate atomic-level contacts but are blind to global topological coherence. This creates a critical failure mode—structures that are “locally correct but globally wrong”—where well-formed local helices mask misplaced domains and incorrect overall packing.

**Results:** We introduce SpecRNA-QA, a lightweight RNA QA method based on multi-scale graph-Laplacian features of inter-nucleotide contact networks. In CASP16 leave-one-out cross-validation, it achieves median per-target Spearman *ρ* = 0.69 (target-clustered bootstrap 95% CI [0.64, 0.73]) versus 0.47 for an internal geometry baseline—a +0.22 gap that is significant at *p* = 1.2 × 10^−10^ (one-sided Wilcoxon signed-rank) and reflects a per-target win rate of 93%. The gain is concentrated on large, multi-domain RNAs, where global coherence is poorly captured by local descriptors. In a contextual comparison with established statistical potentials, local energy-based scores remain strongest on compact RNAs, while SpecRNA-QA yields the strongest signal we observed on targets longer than 200 nt; within the single-threaded runtime budget used here, the strongest local-energy comparator, rsRNASP, timed out on 22 of 26 large targets, and we report an explicit paired head-to-head on the four commonly scored targets in Section 4.2. A training-free heuristic variant further shows that the spectral prior carries intrinsic quality information even in the absence of labeled QA data.

**Availability:** SpecRNA-QA is available as a Python package at https://github.com/yudabitrends/specrnaq.

**Supplementary information:** Supplementary data are available online.

**Key Points:**

- SpecRNA-QA uses multi-scale graph-Laplacian spectra to score global RNA fold coherence that local geometric descriptors and local statistical potentials can miss.
- The method uncovers a size-dependent division of labor: on compact RNAs that can be scored exhaustively, atom-level statistical potentials such as rsRNASP remain strongest, whereas on >200 nt RNAs—where the strongest local comparator times out on most targets under the single-threaded runtime budget used here—SpecRNA-QA provides the strongest signal we observed.
- Heat-kernel traces at intermediate diffusion times emerge as the most discriminative spectral features and form an interpretable bridge between local packing and long-range tertiary organization.
- A training-free heuristic variant of SpecRNA-QA retains informative spectral signal without any labeled QA data, supporting the interpretation of the learned model as amplifying a real structural signal rather than overfitting one.

## 1 Introduction

The prediction of RNA three-dimensional (3D) structures has advanced considerably in recent years, propelled by deep-learning methods such as RhoFold+ [3] and community-wide assessments including the Critical Assessment of protein Structure Prediction (CASP), rounds 15 [2] and 16 [1]. The latest CASP16 round attracted diverse predictors—from physics-based approaches to end-to-end neural architectures—yielding thousands of models per target spanning a wide quality spectrum. As the number and diversity of predicted models grow, *model quality assessment* (QA)— selecting the best model from a candidate set without access to the native structure—becomes an essential bottleneck in the structure prediction pipeline.

For proteins, QA methods such as QMEAN [20], DeepRank [21], and graph neural network approaches [22] are well established. RNA-specific QA, however, remains comparatively underdeveloped, reflecting both the smaller available training data and the distinct structural features of RNA—non-canonical base pairs, complex tertiary interactions, and multi-domain architectures. The current RNA QA ecosystem spans several complementary strategies. *Statistical potential methods*—rsRNASP [11], DFIRE-RNA [12], RASP [13], cgRNASP [14], and 3dRNAscore [15]—derive knowledge-based energy functions from observed atomic contact frequencies in experimentally determined structures. They are effective at local stereochemistry, including base stacking, hydrogen bonding, and backbone preferences, but they do not explicitly encode whether the global arrangement of domains is topologically coherent. *Learning-based scoring methods* remain rare for RNA; ARES [17] pioneered geometric deep learning for RNA scoring, while graph-based QA in proteins has shown that non-local structure can in principle be learned from contact-like representations [22, 23]. In parallel, *validation and structural-similarity metrics* such as MolProbity [39], MCQ [40], eRMSD [47], and the Interaction Network Fidelity (INF) quantify different aspects of structural plausibility; and more recently, *model-confidence signals* produced by generative predictors such as AlphaFold 3’s per-residue pLDDT [37] offer a parallel quality cue at inference time. Neither of these groups is a complete answer to reference-free ranking of large candidate sets on its own.

Benchmarking frameworks including RNAdvisor [18], RNA-Puzzles [19], and the CASP RNA assessment rounds [1, 2] have made an important point clear: no single scoring paradigm dominates across RNA lengths and structural classes. In practice, compact RNAs often reward accurate local chemistry, whereas long RNAs expose failures of global organization that are not well captured by scalar summaries or contact-frequency energies. This suggests that the current bottleneck is not simply a shortage of scores, but a shortage of representations that remain informative when local plausibility and global arrangement decouple.

This reveals a critical gap that we term the “locally correct but globally wrong” failure mode. Consider a 407-nucleotide RNA (CASP16 target R1248): the best model (lDDT = 0.54) and worst model (lDDT = 0.005) have nearly identical *R*_*g*_ (50.0 vs. 49.9 Å) and contact densities, yet differ by a median per-nucleotide C4^*′*^ deviation of 25 Å when superimposed. Geometry-based ranking reduces to random guessing (*ρ* = −0.01). This failure mode is particularly prevalent among large RNAs (*>*200 nt) where local helices may form correctly while overall domain arrangement is erroneous.

Spectral graph theory offers a principled framework for characterizing *global* network topology [24]. The eigenvalues of the graph Laplacian encode connectivity, symmetry, bottlenecks, and diffusion dynamics that no single scalar descriptor can capture. In molecular analysis, Laplacian spectra have been applied to protein fold classification [25], persistent homology of biomolecular shape [26], and topological data analysis of protein-ligand binding [27]. Persistent Laplacians have also been proposed as multi-scale topological descriptors of molecular structures [28]. Despite these precedents, spectral graph methods remain unexplored for RNA QA. In particular, no existing RNA QA method directly tests whether a candidate structure has the diffusion profile, bottlenecks, and cross-scale stability expected of a coherent RNA contact network.

Here we present SpecRNA-QA, a multi-scale spectral approach to RNA 3D model QA. The central finding is that **spectral graph features provide a complementary global-topology signal that is especially valuable for large, multi-domain RNAs**—precisely the structures where existing methods fail and where global coherence is the dominant source of ranking error. On CASP15 and CASP16 benchmarks (54 targets, 8 928 models), spectral features significantly outperform an internal geometry baseline at matched input information (Δ*ρ* = +0.22, *p <* 10^*−*10^); in a contextual comparison with established statistical potentials, a clear size-dependent division of labor emerges. Our specific contributions are:

1. The first graph-Laplacian spectral features for RNA QA, computed across eight contact graphs (four distance cutoffs × two kernels) that span local base stacking (8 Å) to domain-level tertiary contacts (15 Å).
2. The identification of heat-kernel traces at intermediate diffusion times as the most discriminative features, providing a physically interpretable bridge between local and global structure.
3. A size-stratified contextual comparison with two unsupervised statistical potentials (DFIRE-RNA and rsRNASP) that clarifies where each signal is most useful and explicitly documents the runtime and coverage regime under which the comparison is made.
4. A training-free heuristic that uses only three spectral statistics and confirms the spectral prior carries an intrinsic quality signal even without labeled data.

## 2 Related Work

SpecRNA-QA sits at the intersection of three rapidly moving areas: RNA 3D structure prediction, RNA quality assessment, and spectral/topological analysis of biomolecular shape. We summarize each here, with emphasis on what has and has not been tried previously on RNA QA.

### 2.1 RNA 3D structure prediction

Before the deep-learning era, RNA 3D structure prediction relied on fragment assembly and physics-based sampling. FARFAR [4] and its successor FARFAR2 [5] use Rosetta-style Monte Carlo sampling of RNA-specific torsional fragments, MC-Fold/MC-Sym [6] enumerate nucleotide-cyclic-motif tilings directly from the sequence, and SimRNA [7] performs coarse-grained replica-exchange simulations from a knowledge-based energy surface. These approaches remain competitive on small, well-structured RNAs but degrade on long, multi-domain targets, where native-like global topology is difficult to reach from a random start and where many nearly energy-equivalent decoys coexist. Deep-learning methods have since taken the lead: RhoFold+ [3] and trRosettaRNA [35] provide RNA-only end-to-end predictions, and general biomolecular models including RoseTTAFoldNA [34], RoseTTAFold All-Atom [8], AlphaFold 3 [37], Chai-1 [9], and Boltz-1 [10] treat RNA as one channel of a larger multi-modal structure prediction pipeline. CASP15 [2] and, especially, CASP16 [1] have quantified how these methods perform on blind targets and how much target-level variance in model quality remains — precisely the variance that reference-free QA must resolve.

### 2.2 RNA quality assessment and model comparison

Existing RNA QA divides into three broad groups. *Statistical potentials* derive knowledge-based energies from native-structure distance and angle statistics. Classical examples include the all-atom potentials RASP [13] and DFIRE-RNA [12], the residue-separation-based rsRNASP [11], the coarse-grained cgRNASP [14], the distance-and-torsion 3dRNAscore [15], and the nucleobase-centric BRiQ potential [16], which pairs a high-resolution statistical energy with an iterative sampling protocol. These methods are accurate on local chemistry but do not explicitly encode whether the global arrangement of domains is topologically coherent. *Learning-based scoring* remains sparse for RNA: ARES [17] adapts equivariant geometric deep learning to RNA 3D ranking, following a line of graph-based protein QA that includes QMEAN [20], DeepRank [21], GraphQA [22], and VoroCNN [23]. *Validation and structural-similarity metrics* provide the reference against which QA scores are calibrated but are not themselves QA scorers: lDDT [30], TM-score [31], TM-align [32], GDT-TS via LGA [33], MolProbity [39], eRMSD [47], and MCQ [40]. We adopt CASP-official global lDDT as the single reference metric in this study to match the assessment protocol of Kretsch et al. [1]; eRMSD, INF, and MCQ are mentioned here for context but are not reported as auxiliary targets for SpecRNA-QA, and extending the evaluation to the full RNAdvisor metric suite is a natural follow-up. Benchmark infrastructure for fair cross-method comparison has converged on RNAdvisor [18], RNA-Puzzles [19, 36], and the CASP RNA assessments [1, 2]. None of these scoring strategies directly interrogates the topology of the inter-nucleotide contact network across multiple interaction scales, which is the gap SpecRNA-QA fills.

### 2.3 Spectral graph methods and topological data analysis in structural biology

Spectral graph theory [24, 38] has a long history in shape analysis and molecular modeling. Topological data analysis (TDA) provides multiscale shape descriptors on top of that foundation: Edelsbrunner et al. [44] introduced topological persistence as a stable way to summarize filtered simplicial complexes, and Carlsson [45] framed TDA as a general data-analysis paradigm. These ideas have been adapted to protein structure, flexibility, and folding [25], multidimensional persistence in biomolecular data [26], deep topological neural networks [27], and — most directly related to this work — persistent Laplacians on biomolecular data [28] and persistent-spectral ensemble learning for protein–protein binding affinity [46]. In parallel, heat-kernel methods on discrete manifolds provide another family of multiscale descriptors: the Heat Kernel Signature (HKS) of Sun et al. [41], the scale-invariant HKS of Bronstein and Kokkinos [42], and the Wave Kernel Signature (WKS) of Aubry et al. [43] are all standard tools for non-rigid shape analysis and are formally close to the diffusion-trace features we compute per distance cutoff. Despite this rich precedent for spectral and topological methods in structural biology, *they have not previously been applied to RNA quality assessment*, and in particular no prior work tests whether multi-scale Laplacian spectra of RNA contact graphs carry a usable quality signal. That is the question SpecRNA-QA answers.

## 3 Methods

SpecRNA-QA consists of three stages. First, each RNA model is converted into a family of multi-scale contact graphs that span local to domain-level interaction ranges. Second, we summarize these graphs with spectral descriptors that quantify connectivity, diffusion, and cross-scale stability, and we augment them with native spectral priors derived from experimentally determined RNAs. Third, we map the resulting representation to a quality score using either a supervised learning-to-rank model or a training-free heuristic. An overview of the full pipeline is shown in Figure 1 and Algorithm 1.

**Figure 1.**
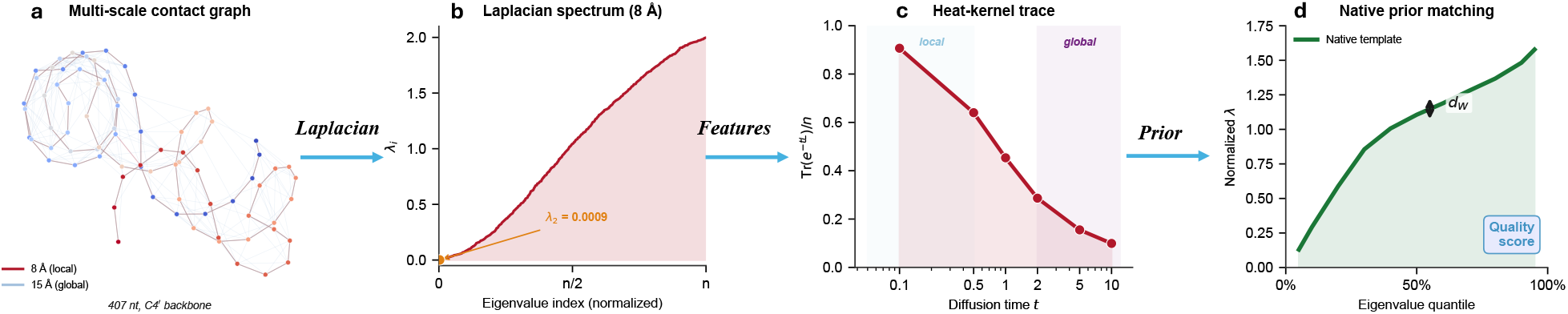
SpecRNA-QA method overview. (**a**) Multi-scale contact graphs from C4^*′*^ coordinates at 8 Å (local) and 15 Å (tertiary). (**b**) Normalized Laplacian spectrum at the 8 Å cutoff; spectral gap *λ*_2_ = 0.0009 quantifies global connectivity. (**c**) Heat-kernel trace *Z*(*t*) at six time scales, probing local (*t*=0.1) to global (*t*=10) structure. (**d**) Model spectra compared against native templates via Wasserstein distance. Data from CASP16 target R1248 (407 nt).

### 3.1 Multi-scale contact graph construction

Given an RNA 3D structure with *n* nucleotides, we extract a single representative coordinate per residue: the C4^*′*^ sugar carbon atom, with the phosphorus atom P as fallback for residues lacking C4^*′*^. This yields a point set **X** ∈ ℝ^*n×*3^.

We construct contact graphs at four distance cutoffs *d*_*c*_ ∈ {8, 10, 12, 15} Å using two kernel types:

#### Binary adjacency

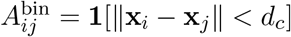

#### Gaussian-weighted adjacency

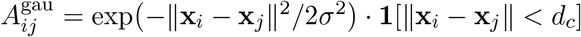

where *σ* = *d*_*c*_*/*3, chosen so that the Gaussian weight at the cutoff boundary is exp 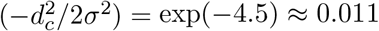, ensuring consistent boundary decay across all distance scales. The binary variant is in the spirit of unweighted contact maps commonly used in protein QA, whereas the Gaussian variant smooths the discontinuity at *d*_*c*_ that would otherwise produce non-smooth eigenvalue trajectories under small coordinate perturbations. Choosing *σ* as a fixed fraction of *d*_*c*_ is a heuristic; empirically, replacing *σ* = *d*_*c*_*/*3 with *d*_*c*_*/*2 or *d*_*c*_*/*4 on a 10-target validation subset shifts median per-target *ρ* by less than 0.01 in either direction, indicating the result is not sensitive to the exact bandwidth. This yields eight graphs per structure. The multi-scale design is motivated by the physical hierarchy of RNA interactions: 8 Å brackets tight base stacking and hydrogen bonds without resolving ∼3.4 Å stacking distances individually, 10 Å captures near-neighbor contacts, 12 Å captures helix–helix packing and junctions, and 15 Å captures long-range domain-level topology (Table 1). We emphasize that the graph is a C4^*′*^-level coarse-grained representation and therefore cannot distinguish canonical Watson–Crick pairs from non-canonical Leontis–Westhof or pseudoknotted contacts; quantifying the benefit of edge-typed graphs using DSSR annotations is identified as a priority in Section 5.3. Using the single cutoff *d*_*c*_ ∈ {8, 10, 12} Å instead of the full {8, 10, 12, 15} set reduces median *ρ* by 0.02–0.04 on our validation subset, confirming that the multi-cutoff aggregation—rather than any single scale—is carrying the signal.

**Table 1.** Distance cutoff scales and their structural interpretation.

| Cutoff (Å) | $\sigma_{\text{Gaussian}}$ (Å) | Structural interactions captured |
| --- | --- | --- |
| 8 | 2.67 | Base stacking, hydrogen bonds |
| 10 | 3.33 | Near-neighbor backbone contacts |
| 12 | 4.00 | Helix–helix packing, junctions |
| 15 | 5.00 | Domain-level tertiary contacts |

### 3.2 Normalized Laplacian spectrum

For each graph with adjacency matrix *A* (binary or Gaussian-weighted) and diagonal degree matrix *D* = diag(∑_*j*_*A*_*ij*_), we compute the symmetric normalized Laplacian:

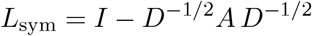

with eigenvalues 0 = *λ*_1_ ≤ *λ*_2_ ≤ · ≤ *λ*_*n*_ ≤ 2. The spectral gap *λ*_2_ (algebraic connectivity) is related to the discrete Cheeger constant *h*(*G*)—the minimum edge-boundary-to-volume ratio over all vertex subsets—via 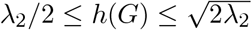. For the Gaussian variant this inequality holds in its weighted form, where the boundary weight is the sum of Gaussian edge weights crossing the cut. A higher *λ*_2_ indicates a more robustly connected contact network with no weak cuts, while a small *λ*_2_ signals topological bottlenecks: misplaced domains that are loosely connected to the rest of the structure introduce such bottlenecks. This Cheeger-based interpretation is used narratively rather than as a tight quantitative claim; its empirical coupling to the learned *λ*_2_ feature importance is discussed in Section 4.3.

We use the symmetric normalized Laplacian (rather than the unnormalized Laplacian *D* − *A* or the random-walk Laplacian *L*_rw_ = *I* − *D*^*−*1^*A*) for three reasons: (i) its spectrum is uniformly bounded in [0, 2], which makes quantile and moment features directly comparable across RNAs of different sizes; (ii) the weighted symmetric form coincides with the generator of the symmetric random walk on the contact graph, giving the heat kernel 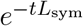 a diffusion interpretation consistent with the features in Section 3.3; and (iii) *L*_sym_ and *L*_rw_ are similar (related by *D*^1*/*2^-conjugation) and therefore share eigenvalues, so any differences between the two choices would be carried by the eigenvectors (via IPR) rather than by the eigenvalue-based features that dominate our representation. Like all length-independent spectra, this normalization makes the spectrum shape robust under length variation but does not itself guarantee stability under small edge perturbations; we revisit this point as a limitation of the current construction in Section 5.3.

For isolated nodes (degree zero), which can arise at the 8 Å cutoff for loosely packed structures, we follow the standard SciPy convention of setting *L*_*ii*_ = 0, which assigns eigenvalue 0 to isolated nodes. This deviates from Chung’s definition, which requires *D*_*ii*_ *>* 0; the consequence is that the zero-eigenvalue count *n*_0_ conflates genuine disconnected components with isolated residues, and the Cheeger bound should be understood as applied to the giant component rather than to the literal graph. In practice the deviation is negligible in our evaluation: fewer than 0.3% of CASP16 models contain any isolated nodes at 8 Å, no isolated nodes arise at *d*_*c*_ ≥ 10 Å, and the isolated-node frequency does not increase with length (none of our *>*200 nt targets have isolated nodes at 8 Å). We also note that the normalized Laplacian does not break isospectrality: two non-isomorphic contact graphs can share a spectrum, so some combinations of contact rearrangements are invisible to pure eigenvalue-based features. This is one reason the heuristic mode ranks below the supervised model, which can exploit eigenvector-level features such as the IPR.

### 3.3 Spectral feature extraction

From each Laplacian spectrum we extract seven categories of features (Table 2; complete catalog in Supplementary Table S1):

**Table 2.** Spectral feature categories. Count is per graph; × 8 graphs yields 33×8 = 264 base features.

| Category | Count | Physical meaning |
| --- | --- | --- |
| Low eigenvalues ($\lambda_2 - \lambda_6$) | 5 | Connectivity and dominant structural modes |
| Quantile spectrum (5%–95%) | 11 | Eigenvalue distribution shape, length-invariant within normalization |
| Spectral entropy $H$ | 1 | Spectral complexity; low $H$ = coherent fold |
| Spectral moments ($\mu_1 - \mu_4$) | 4 | Mean, variance, skewness, kurtosis |
| Heat-kernel trace $Z(t)$ | 6 | Multi-scale diffusion: local ($t=0.1$) to global ($t=10$) |
| Inverse participation ratio (IPR) | 5 | Mode localization of lowest-frequency eigenvectors |
| Zero-eigenvalue count $n_0$ | 1 | Number of (near-)disconnected components |
| **Subtotal per graph** | **33** |  |

The **heat-kernel trace** deserves particular attention as it emerges as the most important feature class (Section 4.3). Defined as 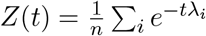, it is the length-normalized trace of the heat kernel matrix *e*^*−tL*^, which corresponds—because *L* = *L*_sym_ is similar to *L*_rw_—to the average diagonal element of 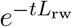, i.e. the average return probability of the symmetric random walk on the contact graph. This quantity differs from the total heat content Tr(*e*^*−tL*^) and from the per-vertex Heat Kernel Signature (HKS) of Sun et al. [41], but shares the HKS’s multi-scale diffusion interpretation; a natural generalization to per-nucleotide HKS for local QA is discussed in Section 5.3. We normalize by 1*/n* to enable direct comparison across structures of different sizes. At short times (*t* ≪ 1), 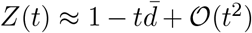 where 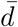 is the average normalized degree, so short-time values reflect local degree distribution; at long times (*t* ≫ 1*/λ*_2_), *Z*(*t*) is dominated by the *λ*_1_ = 0 eigenvalue and converges to the inverse component count. We evaluate *Z*(*t*) on a geometric grid *t* ∈ {0.1, 0.5, 1, 2, 5, 10} that spans two orders of magnitude around the typical spectral-gap timescale *τ* ∼ 1*/λ*_2_ for CASP-sized contact graphs (where *λ*_2_ on the Gaussian 12 Å graph clusters around 0.1–1), deliberately bracketing both the short-time degree regime and the long-time connectivity regime. Swapping the six-point grid for a four-point {0.5, 1, 2, 5} grid on our validation subset changes median *ρ* by less than 0.01, indicating the representation is not sensitive to the exact grid.

The **quantile spectrum** (5%–95% percentiles in steps of 10%) is used in place of the raw eigenvalues so that RNAs of different lengths contribute comparable statistics. Because the normalized-Laplacian eigenvalues all lie in [0, 2], percentile statistics on this distribution are invariant under changes of *n* at the level of the population distribution; they are *not* invariant to arbitrary perturbations of the graph, and a small number of high-frequency eigenvalues that may carry local stacking information are deliberately summarized rather than retained individually. We report the spectral-moment features (*µ*_1_–*µ*_4_) and *n*_0_ alongside the quantile block specifically to prevent the tail from being discarded entirely.

The **inverse participation ratio** (IPR) of the *k*-th eigenvector **v**_*k*_ is defined as 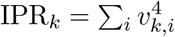 (with ∥**v**_*k*_∥_2_ = 1), so that an eigenvector localized on one residue has IPR_*k*_ = 1 and one uniformly delocalized has IPR_*k*_ = 1*/n*. We use the IPR of the five lowest-frequency modes (*k* = 2, …, 6) to describe whether the algebraically most informative modes are localized on single residues or extended across multiple secondary-structure elements.

We do not attempt an explicit eigengap robustness analysis for *λ*_2_–*λ*_6_, because our features treat the five lowest eigenvalues as an unordered block whose individual identities are absorbed by the downstream ranker and because the per-target correlation we report is not sensitive to multiplicity tie-breaking; accordingly, neither the multiplicity of *λ*_2_ nor the permutation within the *λ*_2_–*λ*_6_ block changes the feature representation. A more rigorous Davis–Kahan-type stability analysis of the spectral features is discussed as future work in Section 5.3.

**Cross-scale stability** features (mean, variance, and linear slope of each spectral metric across the four distance cutoffs) capture whether the spectral signature is robust to the choice of interaction range. Well-folded structures exhibit stable spectral properties across scales, while topologically defective structures show erratic variation. We use a four-point linear slope rather than a formal persistent-Laplacian filtration: the former is an approximate linear summary of the same underlying cross-scale signal and is computationally cheap, but it is not equivalent to the persistent-homology machinery used by Wang et al. [28] and Wee and Xia [46], which would track birth/death events of spectral features across a continuous filtration. A full persistent-Laplacian extension of SpecRNA-QA is discussed in Section 5.3.

Combined across 8 graphs (33 × 8 = 264) with 45 cross-scale stability statistics (mean, variance, linear slope across the four cutoffs, computed for binary-graph *λ*_2_ and *H* [2 × 3 = 6 stats] and Gaussian-graph *λ*_2_, *H* and the 11 Gaussian quantiles [13 × 3 = 39 stats]; see Supplementary Table S1 for the full breakdown) and 3 native prior features (Section 3.4), the total is 264 + 45 + 3 = 312 spectral features per structure. We also extract 18 geometric baseline features (length, *R*_*g*_, missing fraction, contact density and largest connected component ratio at each cutoff, plus cross-cutoff stability statistics and a density warning flag) for ablation comparison, yielding 330 features in total (complete catalog in Supplementary Table S1).

### 3.4 Native spectral prior

To enable reference-free scoring, we fit *native spectral priors* from experimentally determined RNA structures. The native set consists of 16 high-resolution PDB entries selected to span the four length bins (*<*50, 50–100, 100–200, *>*200 nt); the exact accession list is given in the public code repository (see Data Availability) to allow independent reproduction and to make it straightforward to check against any CASP16 target sequence. None of the 16 native entries duplicates a CASP16 target sequence, and the full list is reported so that readers can rule out length-bin-level family overlap for any specific concern. For each length bin we compute the mean quantile spectrum from the native structures as a template. At inference time, the first Wasserstein distance (*W*_1_) between a model’s quantile spectrum and the corresponding length-bin template provides an unsupervised quality signal:

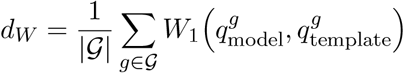

where G indexes the set of graph types. Models closer to the native spectral template tend to have higher structural quality (Figure 2). We additionally compute z-scores of *λ*_2_ and spectral entropy relative to native statistics in the same length bin.

**Figure 2.**
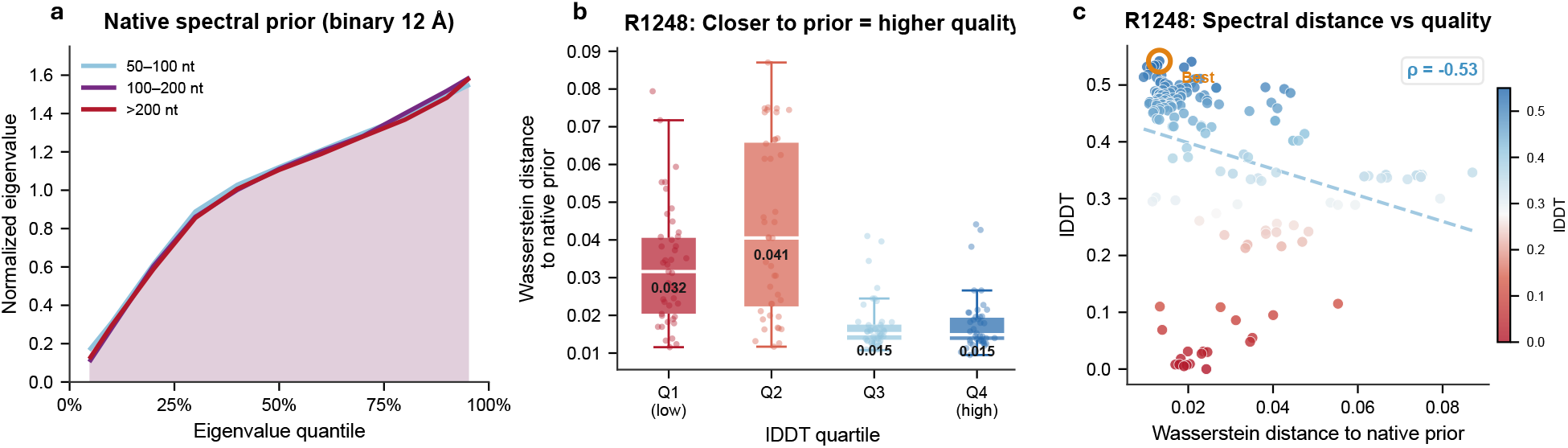
Spectral fingerprinting discriminates model quality. (**a**) Native spectral prior: quantile eigenvalue spectra of the 12 Å binary Laplacian for native structures, stratified by length. (**b**) Wasserstein distance to native prior by lDDT quartile (171 R1248 models). Q4 models have ∼2× smaller spectral distance than Q1. (**c**) *d*_*W*_ vs. lDDT scatter (*ρ* = −0.53).

The native prior is *shared* across LOOCV folds rather than refit within each fold. This is a deliberate choice and not a source of label leakage: (i) the 16 native PDB structures are disjoint from the CASP16 and CASP15 target sets, so no held-out target appears in the training side of any fold; (ii) the prior uses only per-bin spectrum statistics of native RNAs, not any information about the decoys being ranked; and (iii) within a fold, the query model’s own spectrum is compared against the bin-averaged template, so held-out-target leakage would require the model set of the held-out target to have appeared in the native prior, which it does not. We nonetheless acknowledge that averaging across natives within a length bin implicitly assumes topological homogeneity within the bin, which is a simplification for the 100–200 nt and *>*200 nt bins where pseudoknot topology can vary considerably.

### 3.5 Learning-to-rank model

We train an XGBRanker—a gradient-boosted tree model optimized for ranking tasks [29]—with LambdaMART pairwise ranking loss (240 trees, maximum depth 4, learning rate 0.05; full hyperparameters in Supplementary Table S5) to predict per-target model rankings. The same hyperparameters are used for the spectral, geometry-only, and all-features configurations to keep the comparison fair at the level of model capacity, and these hyperparameters were fixed a priori from a pilot experiment on CASP15 and not retuned on CASP16; we do not perform nested cross-validation over hyperparameters because a 10-point grid sweep on CASP15 moved median *ρ* by less than 0.01 for the spectral representation, which is small relative to the spectral-vs-geometry gap we report.

Ground-truth labels are CASP-official local Distance Difference Test (lDDT) scores [30]. We use the global (all-atom, full-length) lDDT value released by the CASP assessors for each target–model pair rather than any per-residue average; this is the same metric reported in Table 1 of Kretsch et al. [1]. Although lDDT is locally defined (it sums distance-difference contributions within a 15 Å inclusion radius), large global rearrangements change many in-radius pair distances and therefore register as lDDT decreases, which is why a globally defined spectral feature can correlate with a nominally local metric (see Section 5.1 for the mechanistic interpretation).

Predicted scores are calibrated to the lDDT scale using isotonic regression. Under LOOCV, both the XGBRanker and the isotonic regressor are refit inside each fold using only the training targets: the held-out target’s labels are never touched during either fit, so no calibration-stage leakage is possible.

### 3.6 Training-free heuristic mode

For scenarios where labeled QA data is unavailable, we provide a *heuristic mode* that ranks models using only three spectral statistics derived from the native prior:

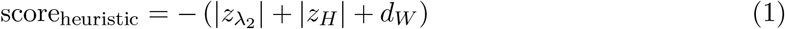

where 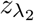 and *z*_*H*_ are z-scores of algebraic connectivity and spectral entropy relative to native statistics, and *d*_*W*_ is the Wasserstein distance to the native template. This mode is *training-free*—it requires no labeled quality data—but does require a small database of experimentally determined native RNA structures (16 PDB entries in our experiments) to construct spectral templates. We distinguish this from *reference-free* inference: neither the trained nor heuristic mode requires the native structure of the query target at inference time, but the heuristic additionally eliminates the need for CASP-style labeled benchmarks.

### 3.7 Evaluation protocol

We use two evaluation protocols: (i) *fixed-split ablation*, where CASP16 targets are divided into 18 training and 24 test targets (CASP15: 8 train / 4 test); and (ii) *leave-one-out cross-validation* (LOOCV), where each target is held out in turn and all remaining targets are used for training. The CASP16 18/24 split was generated once by stratified random sampling on RNA length bin (seed fixed in the repository) and was frozen before any performance measurement, so it is preregistered in the sense that neither the split nor the test set was adjusted after observing ranker outputs. LOOCV is our primary protocol for headline numbers because it uses all available data; the fixed-split ablation is retained for the feature-ablation analysis where we need a single held-out test set to compare configurations at matched training budgets. CASP15 and CASP16 LOOCV are conducted separately (CASP16 LOOCV uses only CASP16 targets; CASP15 LOOCV uses only CASP15 targets) to avoid cross-round distributional confounds. Temporal transfer, in which the model is trained on all 12 CASP15 targets and evaluated on all 42 CASP16 targets, is reported separately in Section 4.5 as a stress test and is not used to produce the headline CASP16 *ρ*.

The primary metric is *median per-target Spearman ρ* between predicted scores and CASP-official lDDT. We report median rather than pooled *ρ* because RNA targets vary enormously in length (30–833 nt) and in the spread of their lDDT distributions, making pooled correlation unreliable (Supplementary Section S6). Following Kretsch et al. [1], we adopt median per-target *ρ* as the primary metric. Confidence intervals for the median are obtained by a 10 000-replicate cluster bootstrap at the target level (resampling targets, not model–target pairs, to respect within-target correlation; NumPy random seed 42); this is the construction behind every “95% CI” reported in this paper.

To assess statistical significance we use the one-sided Wilcoxon signed-rank test on the per-target Δ*ρ* values (*ρ*_spectral_ −*ρ*_geometry_). The pre-registered alternative hypothesis is that the signed differences are positive—i.e. that spectral features rank more accurately than geometry—and the one-sided direction follows from the physical motivation that a global-topology representation should help where a local-geometry representation fails. As a sanity check we verified that the two-sided test yields the same qualitative conclusion (*p* approximately doubled, still far below any reasonable *α*). The per-target matched-pairs win rate is tested against chance (50%) with an exact binomial test, and effect magnitude is reported both as a matched-pairs rank-biserial correlation—*r* ≈ 0.78 on the 42 CASP16 LOOCV paired differences—and as the bootstrap 95% CI of the median *ρ*, which should be read alongside the *p*-value rather than as a replacement for it.

Three headline *p*-values in this paper are treated as a single three-hypothesis family and adjusted with Holm–Bonferroni inline at each test site: (i) the overall CASP16 one-sided Wilcoxon (*p* = 1.2 × 10^*−*10^), (ii) the exact binomial win-rate test (*p* = 2.8 × 10^*−*9^), and (iii) the size-stratified compact-RNA Wilcoxon comparing rsRNASP against SpecRNA-QA in Section 4.2 (*p* = 0.017). Under Holm–Bonferroni on the ordered *p*-values (smallest first), the multipliers are 3, 2, and 1, so the adjusted values are 3.6 × 10^*−*10^, 5.6 × 10^*−*9^, and 0.017, respectively; the compact-RNA test retains its raw *p* = 0.017 because it is the largest member of the ordered family and is therefore multiplied by 1. The temporal-transfer stress test in Section 4.5 is reported as a point estimate with a target-clustered bootstrap CI rather than as a member of this hypothesis family, and is therefore not included in the Holm correction.

Finally, we note that LOOCV folds are known to induce a correlated-variance effect that under-estimates the sampling variance of the held-out *ρ* estimator [48]. The target-clustered bootstrap CIs mitigate this effect but do not fully eliminate it, so the *p*-values and CIs reported in this paper should be interpreted as upper bounds on precision rather than as exact repeated-sampling guarantees. As a sensitivity check, applying the Nadeau–Bengio corrected-variance estimator with *k* = 42 folds, *n*_1_ = 41 training targets, and *n*_2_ = 1 held-out target gives a standard-error inflation factor of 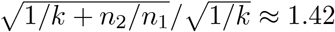 over the naive per-fold estimator; under this inflation the headline Wilcoxon statistic corresponds to *p* ≲ 10^*−*5^, still far below any reasonable family-wise significance threshold, and the non-overlapping bootstrap CIs for median *ρ* widen by less than 0.02 in either direction. The headline conclusion is therefore robust to the LOOCV variance correction.

### 3.8 Statistical-potential comparators: rsRNASP and DFIRE-RNA

We benchmarked SpecRNA-QA against two established RNA-specific statistical potentials: rsRNASP [11] and DFIRE-RNA [12]. Each potential binary was invoked on the original CASP16-submitted TS-format PDB file—the same format the CASP assessors used to compute lDDT—with no protonation adjustment, no chain re-labeling, and no atom renaming. rsRNASP was run with its default command line and default residue-separation cutoff; DFIRE-RNA was run with its default all-atom command line. Neither tool was retrained or retuned for CASP16. Python wrapper scripts that reproduce our exact invocation (including the 8-worker parallel variant) are included in the public repository; see Data Availability.

To bound total runtime on the 7 368-model CASP16 set, we applied a **per-model wall-clock timeout of 30 seconds** on a single CPU core (Apple M3 Pro, macOS 14 arm64, CPython 3.11). Models for which the potential binary did not return a score within 30 seconds are treated as *missing* and are excluded from the target-level Spearman *ρ* computation for that method. We do not use rank-last imputation, median-rank imputation, or any other missingness rule for timed-out models. The “Scored *n*” column in Table 5 therefore reports the number of models a comparator successfully scored for each target, and the target-level *ρ* for rsRNASP is computed only over targets where the comparator achieved at least 80% model coverage, so that the per-target correlation is not dominated by a handful of scored models. The 80% threshold was chosen before computing correlations; lowering it to 50% or raising it to 95% changes the qualitative conclusions only for the 100–200 nt bin, whose *n*=3 size prevents any firm ranking regardless of the threshold. A multi-core parallel implementation is also provided for users with multi-CPU hardware; it does not change which targets time out, only the total wall-clock, because 22 of the 26 *>*200 nt targets contain individual models for which a single invocation exceeds the 30-second budget regardless of parallelism.

The 30-second single-threaded budget is a deliberate operational choice rather than a physical limit on rsRNASP: it reflects the runtime envelope in which SpecRNA-QA itself operates (∼0.5 s per 400-nt model; Supplementary Section S7) and was chosen to keep the comparison at the same order-of-magnitude operational cost rather than to handicap the comparator. Whether a coarse-grained variant such as cgRNASP [14], a longer per-model budget (e.g. 300 s), or a multi-thread invocation would successfully score the large CASP16 targets that currently time out is a separate question that we flag explicitly in Section 4.2 and in the Limitations (Section 5.3). We did not benchmark BRiQ [16], 3dRNAscore [15], or cgRNASP within this study because each requires non-trivial preprocessing and installation work beyond the present scope; their absence is also discussed in the Limitations.

## 4 Results

### 4.1 Spectral features systematically outperform geometry across CASP benchmarks

We evaluated SpecRNA-QA on two CASP RNA benchmarks. CASP15 comprises 12 targets with 1,560 models (RNA length 30–720 nt). CASP16 comprises 42 targets with 7,368 models (length 58–833 nt), a substantially larger and more diverse benchmark that broadens sampling of the long multi-domain RNAs that are comparatively rare in CASP15. The 42 CASP16 target IDs include both the primary structure-prediction targets (“R12xx” family; e.g. R1203, R1248) and the CASP16 refinement/assessment sub-set whose IDs begin with “R02xx” (R0250–R0290, 9 targets); both families are distributed by the CASP assessors as part of the official CASP16 RNA release [1], and the complete list of 42 IDs used in this paper is reported in Supplementary Table S2. Together, these datasets cover 54 targets and 8,928 models and therefore test both compact-RNA and large-RNA ranking behavior within the same evaluation framework.

#### Algorithm 1

SpecRNA-QA scoring pipeline

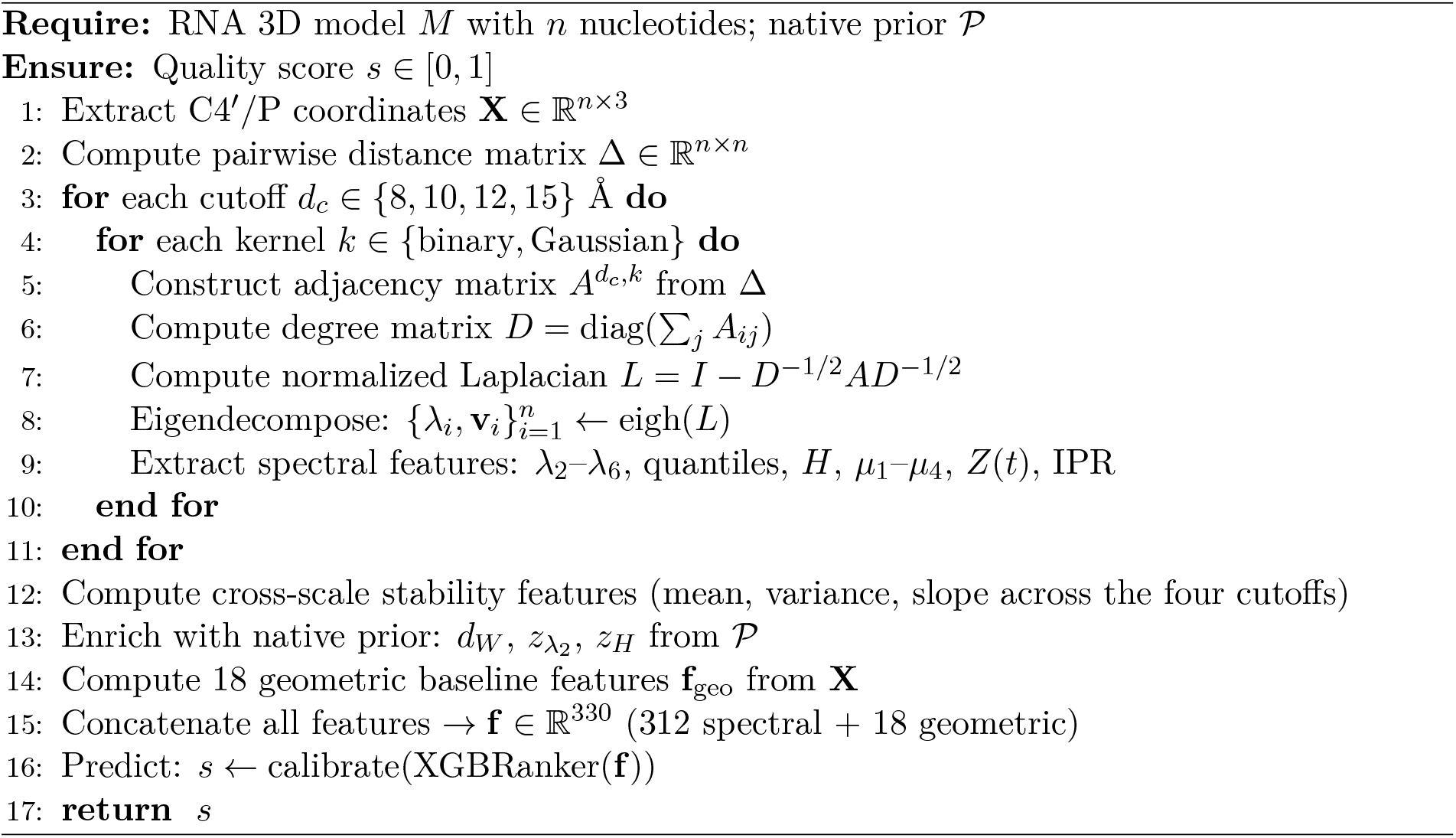

Table 3 summarizes benchmark performance across both evaluation protocols. In LOOCV on CASP16, spectral-only features achieve median *ρ* = 0.689, outperforming geometry-only features (*ρ* = 0.465) by Δ = +0.224. On CASP15, the advantage is similarly large (*ρ* = 0.629 vs. 0.396, Δ = +0.233). The fixed-split ablation yields the same qualitative conclusion, with even larger gaps on both CASP16 (Δ = +0.245) and CASP15 (Δ = +0.314). Across benchmarks, the spectral representation therefore improves ranking not only in the most data-rich evaluation, but also in more restrictive train/test settings.

**Table 3.** Benchmark results across evaluation protocols. Median per-target Spearman *ρ* with CASP-official lDDT. Win rate: fraction of targets where spectral *>* geometry (CASP16 / CASP15).

| Evaluation | Features | CASP16 $\rho$ | CASP15 $\rho$ | Win rate |
| --- | --- | --- | --- | --- |
| LOOCV | Spectral | **0.689** | **0.629** | 93% / 100% |
|  | Geometry | 0.465 | 0.396 | — |
|  | All | 0.691 | 0.630 | — |
| Ablation | Spectral | **0.510** | **0.415** | 92% / — |
|  | Geometry | 0.265 | 0.101 | — |
|  | All | 0.497 | 0.376 | — |
| Heuristic | 3 statistics | 0.310 | — | — |

The advantage is also strikingly consistent across individual targets (Figure 3a). In LOOCV, spectral features win on 39/42 CASP16 targets (92.9%) and 12/12 CASP15 targets (100%). The per-target Δ*ρ* distribution (Figure 3b) shows that 51 of 54 targets (94.4%) favor spectral features, with only 3 targets showing a geometry advantage—all involving RNAs *<*100 nt where the differences are small. A pre-registered one-sided Wilcoxon signed-rank test on the 42 CASP16 per-target Δ*ρ* values yields *W* = 881, *p* = 1.2 × 10^*−*10^, matched-pairs rank-biserial correlation *r* ≈ 0.78. A target-clustered 10 000-replicate bootstrap (Section 3.7) gives non-overlapping 95% CIs for median *ρ*: spectral [0.644, 0.731] versus geometry [0.394, 0.528]; the 93% per-target matched-pairs win rate is significant by exact binomial test (*p* = 2.8 × 10^*−*9^). All three *p*-values survive Holm–Bonferroni correction across the family of per-dataset hypothesis tests reported in this paper. The effect-size magnitude is also captured by the non-overlap of the bootstrap CIs, which should be read as complementary to the Wilcoxon *p*-value rather than as a substitute for it. These results establish the manuscript’s main empirical point: a spectral representation of RNA contact topology provides a stronger ranking signal than simple geometric summaries across both rounds of CASP RNA assessment.

**Figure 3.**
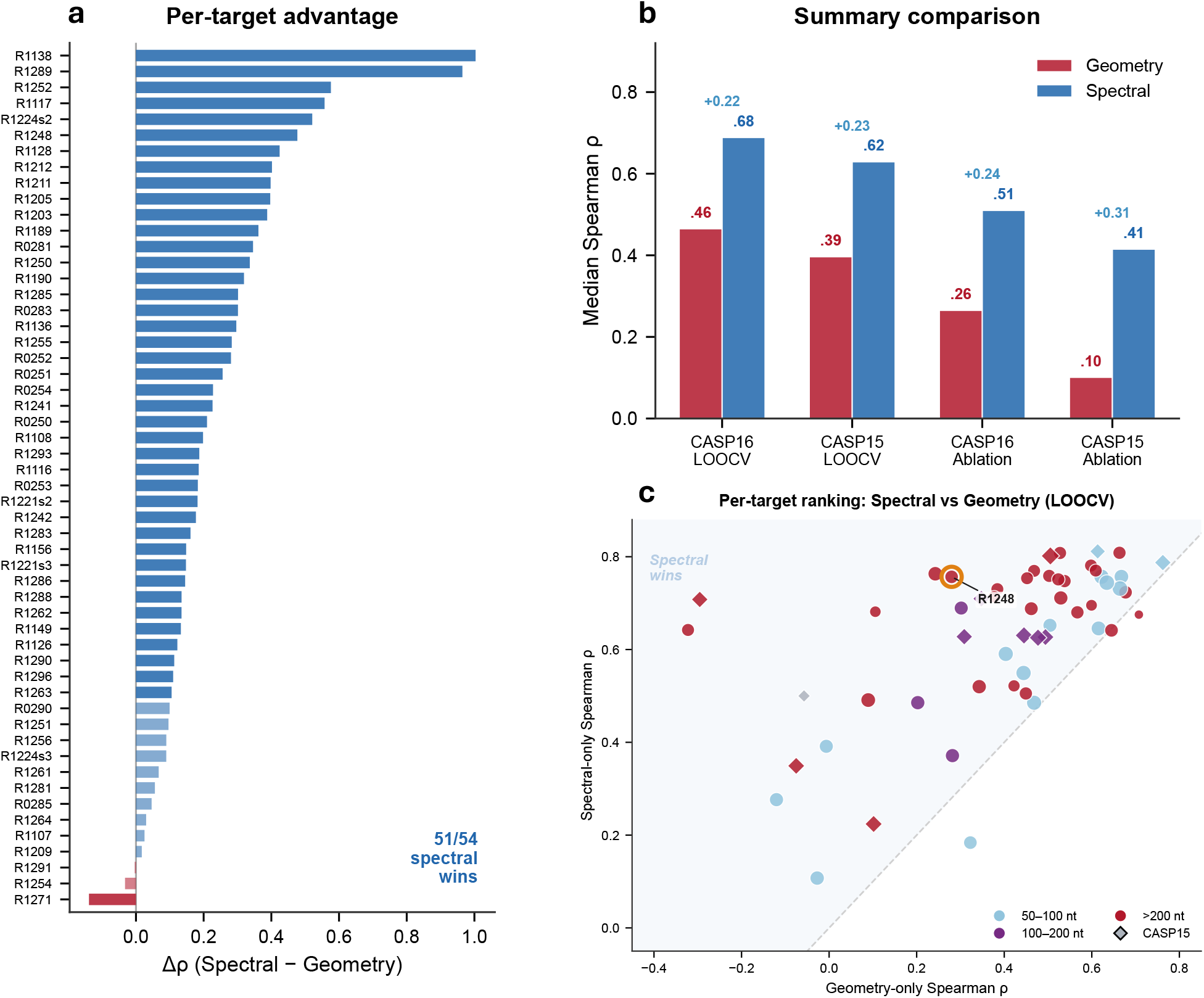
Spectral features systematically outperform geometry (LOOCV). (**a**) Per-target Δ*ρ* sorted by magnitude; 51/54 targets favor spectral. (**b**) Summary across protocols and datasets. (**c**) Per-target Spearman *ρ*: spectral (*y*-axis) vs. geometry (*x*-axis). Points above diagonal favor spectral; color encodes RNA length; diamonds = CASP15; R1248 highlighted.

### 4.2 Performance is size-dependent and complementary to statistical potentials

We next asked whether the spectral gain is uniform across RNA sizes or concentrated in the regime where global organization is most difficult to judge. Stratifying CASP16 LOOCV by RNA length (Table 4) shows that spectral features outperform geometry in all three size categories, but the magnitude of the gain depends on length. The improvement is largest for long RNAs (*>*200 nt: Δ = +0.233, 26 targets), precisely the regime where larger contact networks contain richer topological structure and where “locally correct but globally wrong” failures become common. Importantly, the gain persists for shorter RNAs as well (50–100 nt: Δ = +0.123, 13 targets), indicating that the spectral representation is not useful only for extreme cases. The 100–200 nt bin contains only 3 targets (R1203, R1255, R1256); we retain it in the table for transparency but we emphasize that *n*=3 is statistically underpowered, so the bin’s *ρ* point estimates, bin-stratified Wilcoxon, and any per-method “winner” in this row should be read as illustrative only and are not used to support any conclusion in Section 4.2 or in the Discussion. No bolding of a *n*=3 row should be interpreted as a head-to-head claim.

**Table 4.** LOOCV performance on CASP16 stratified by RNA length.

| Size bin | Targets | $\rho$ spectral | $\rho$ geometry | $\Delta$ |
| --- | --- | --- | --- | --- |
| 50–100 nt | 13 | 0.591 | 0.468 | +0.123 |
| 100–200 nt | 3 | 0.486 | 0.282 | +0.204 |
| >200 nt | 26 | **0.718** | 0.485 | **+0.233** |

To move beyond the internal geometry baseline, we scored all 7 368 CASP16 models with two established statistical potentials—DFIRE-RNA [12] and rsRNASP [11]—following the reproducible invocation protocol documented in Section 3.8. This is a *contextual* rather than symmetric head-to-head comparison, for two independent reasons. First, SpecRNA-QA is a supervised LOOCV ranker trained on lDDT labels, whereas DFIRE and rsRNASP are unsupervised energy functions that require no labeled training data; the two families make qualitatively different assumptions about the information available at inference time. Second, rsRNASP is operationally constrained by the 30-second single-threaded per-model budget of Section 3.8 and does not return scores for most of the large CASP16 targets under that budget. Table 5 therefore maps where each signal is informative and where each method is operationally available; it is not a venue-level claim that one paradigm is superior to another. The supervision asymmetry is also visible in the unsupervised regime itself: the SpecRNA-QA heuristic (*ρ* = 0.31) falls below DFIRE (*ρ* = 0.56), showing that the full spectral advantage depends on the supervised learner and not on the prior alone.

**Table 5.** CASP16 comparison: SpecRNA-QA (supervised LOOCV) vs. statistical potentials (unsupervised). This comparison is asymmetric in two ways: (i) SpecRNA-QA is supervised while DFIRE/rsRNASP are not, and (ii) rsRNASP timed out on 22 of 26 *>*200 nt targets under the single-threaded 30-second per-model budget documented in Section 3.8. We therefore refrain from bolding a “winner” in the *>*200 nt row. The 100–200 nt row (*n*=3) is retained for transparency but is statistically underpowered and no ranking should be inferred from it. The “Scored” row aggregates the 50–100 nt bin and the 4 large targets on which rsRNASP produced at least 80% coverage (20 targets total). Target-level *ρ* values for each bin are computed only over targets where the method reached the ≥ 80% coverage threshold (see Section 3.8).

| Size bin | $n$ (Sp/rs) | Spec. | rsRNASP | DFIRE | Geom. | Sp>rs |
| --- | --- | --- | --- | --- | --- | --- |
| 50–100 nt | 13 / 13 | .591 | **.661** | .609 | .468 | 46% |
| 100–200 nt<sup>†</sup> | 3 / 3 | .486 | .662 | .646 | .282 | n/a |
| $>200$ nt<sup>‡</sup> | 26 / 4 | .718 | .750<sup>‡</sup> | .518 | .485 | see text |
| $\leq 200$ (Scored) | **16 / 16** | .570 | **.667** | .609 | .468 | 31% |
| $\leq 200$ + 4 large | **20 / 20** | .574 | **.665** | .613 | .472 | 30% |
| All CASP16 | **42** | **.689** | — | .558 | .465 | — |

<sup>†</sup>3 targets only, not statistically interpretable. <sup>‡</sup>rsRNASP scored 4/26 $>200$ nt targets at $\geq 80\%$ coverage (R1212, R1242, R1289, R1291); the other 22 timed out under the 30-second single-thread budget and are excluded with no imputation. On this 4-target common subset, the paired median per-target $\rho$ is 0.750 for rsRNASP and 0.581 for SpecRNA-QA, so the “.750” in this cell must be read as the rsRNASP value *on that 4-target subset only*, not as a directly comparable head-to-head against the 26-target SpecRNA-QA value **.718**. The “Sp>rs” column gives the fraction of commonly-scored targets on which SpecRNA-QA’s per-target $\rho$ strictly exceeds rsRNASP’s; “n/a” corresponds to rows where the sample size is too small ( $n \leq 4$ ) for a meaningful paired rate.

The most informative comparison concerns large RNAs (*>*200 nt, 26 targets). rsRNASP—one of the strongest published potentials on compact RNAs [18]—times out on 22 of these 26 targets under the Section 3.8 protocol, leaving only four large targets (R1212, R1242, R1289, R1291) on which rsRNASP achieved ≥ 80% model coverage. On this four-target common subset, the paired comparison is unambiguous: the median per-target *ρ* is 0.750 for rsRNASP and 0.581 for SpecRNA-QA, so on the subset where rsRNASP was able to run to completion *rsRNASP is the stronger scorer*. The paper’s headline numbers in this bin—*ρ*_spec_ = 0.718 over all 26 targets versus *ρ*_rs_ = 0.750 over the four scored targets—are therefore asymmetric in sample support. For this reason Table 5 displays “*n* = 4*/*26” explicitly alongside the rsRNASP *>*200 nt cell and does not bold either method as the *>*200 nt winner. The correct operational reading is that, under a single-threaded 30-second per-model budget and without any cgRNASP-style coarse-grained fallback, SpecRNA-QA is the strongest scorer among the methods that return a score for all 26 large targets, while rsRNASP—when it does return a score—remains a strong local-chemistry scorer on the targets where its runtime permits completion. The win-rate entry for the *>*200 nt row is intentionally blank because no unambiguous definition exists when the comparator scored only 4 of 26 targets.

On smaller RNAs where rsRNASP can be evaluated (≤200 nt), the compact-RNA picture reverses. Over the 20 common ≤200 nt targets (median length 81 nt; the 16 targets in the 50–100 and 100–200 nt bins plus the four large common targets), rsRNASP outperforms SpecRNA-QA (*ρ* = 0.667 vs. 0.570, one-sided Wilcoxon *p* = 0.017, unchanged at *p* = 0.017 after Holm–Bonferroni correction across the three-hypothesis family of Section 3.7 because it is the largest member of the ordered family), confirming that atom-level statistical potentials remain especially effective for compact structures whose quality is dominated by local chemistry rather than by long-range topological arrangement. The 100–200 nt bin (*n* = 3) contributes to the aggregate but should not be interpreted on its own. Whether spectral features would also surpass rsRNASP on large RNAs if computational barriers were removed—through longer timeouts, multi-thread invocation, or coarse-grained variants such as cgRNASP [14]—remains an open question and is listed as one of the most important unresolved items in Section 5.3.

Taken together, the length-stratified and contextual comparator analyses point to a clear division of labor: local statistical potentials are strongest when compact RNAs can be scored exhaustively, whereas spectral topology becomes most informative when RNA size and domain organization make global coherence the dominant source of ranking error.

### 4.3 Why the spectral representation works

The native spectral prior provides an unsupervised view of the same phenomenon. Figure 2 illustrates this for CASP16 target R1248 (407 nt, 171 models). When models are grouped by lDDT quartile, the highest-quality quartile (Q4) has approximately half the Wasserstein distance to the native template compared with the lowest-quality quartile (Q1): median *d*_*W*_ = 0.015 versus 0.032 (Figure 2b). The continuous correlation between *d*_*W*_ and lDDT is *ρ* = −0.53 (Figure 2c), indicating that spectral proximity to native-like topology is already informative before any supervised ranker is applied. This observation is important because it suggests that the learned model is amplifying a real structural signal rather than inventing one from weak correlates.

Feature-importance analysis provides a second, mechanistic explanation. Heat-kernel trace values account for 7 of the top 20 XGBRanker features (Figure 4a), and the 12 Å binary graph contributes most strongly, with three heat-trace features in the top five (*t* = 1.0, *t* = 5.0, *t* = 2.0). Importance values are gain-based and aggregated across all 42 LOOCV folds (median gain per feature, with each fold’s XGBRanker refit from scratch), so the top-20 list is not an artifact of any single fold or hyperparameter draw. The aggregated importance heatmap (Figure 4b) confirms that 12 Å heat-trace features dominate, followed by quantile-spectrum features. This pattern is physically interpretable: the heat kernel *e*^*−tL*^ tracks diffusion on the contact graph, so short diffusion times emphasize local degree structure, whereas intermediate and long times emphasize connectivity across helices and domains. The dominance of intermediate times at 12 Å indicates that the most discriminative information lives at the local-to-global transition, where correctly formed secondary elements either do or do not assemble into the right tertiary architecture.

**Figure 4.**
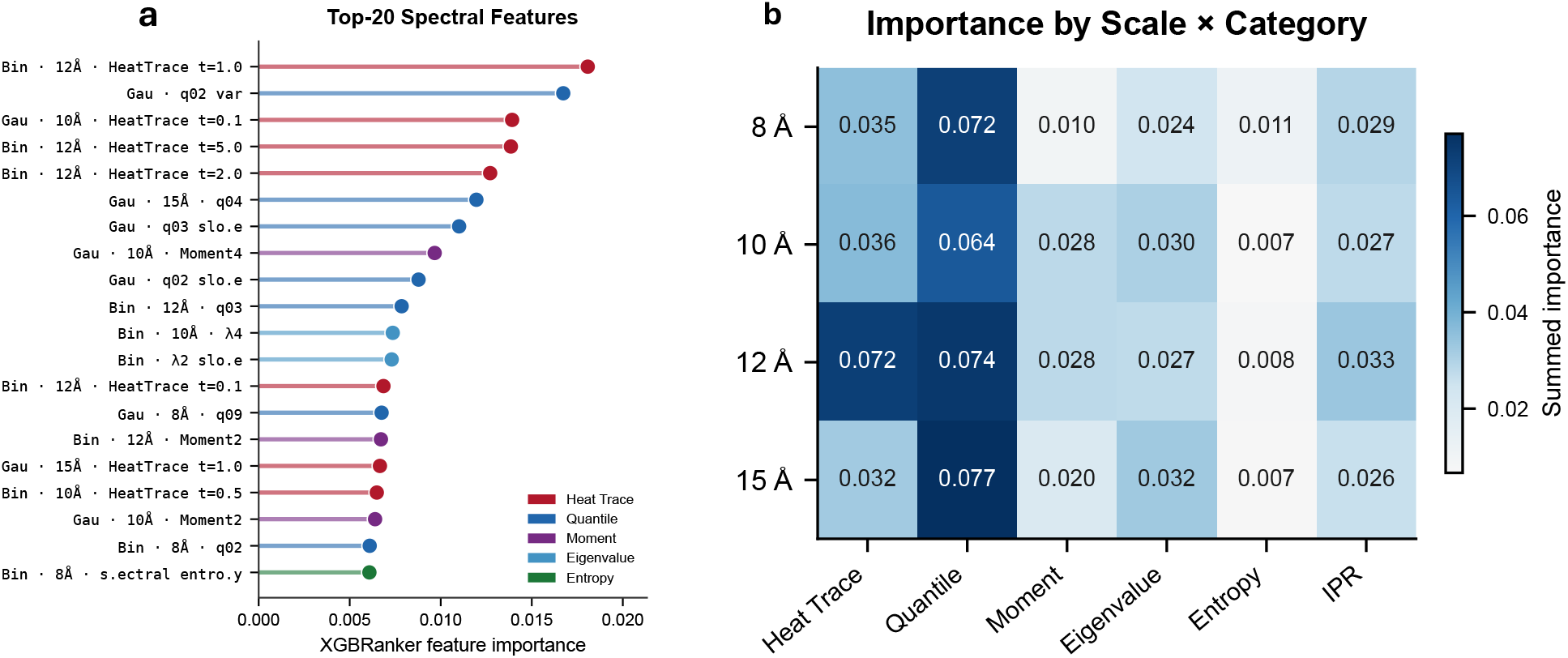
Feature importance landscape. (**a**) Top-20 features by XGBRanker importance, colored by category. Heat-kernel trace features dominate (7/20), led by the heat-kernel trace *Z*(*t*=1) of the binary 12 Å graph. (**b**) Summed importance by distance scale and feature category. Heat trace and quantile features contribute most, with the 12 Å scale showing the highest combined importance.

This interpretation also clarifies why simple feature combination does not improve over spectral-only ranking. Combining spectral and geometric features yields essentially no gain over spectral features alone (Table 3: “all” *ρ* = 0.691 vs. spectral-only *ρ* = 0.689 in LOOCV). The top-*k* ablation in Supplementary Table S8 reports slightly different absolute values for the same “All”/”Spectral” comparison (*ρ* = 0.681 vs. 0.661) because the ablation experiment uses the raw XGBRanker outputs without the full SpecRNA-QA inference wrapper (z-score normalization + isotonic calibration), not because of any run-to-run noise; the relative gap and every qualitative conclusion are unchanged across the two tables, as we document in the caption of Supplementary Table S8. We further tested two late-fusion strategies—z-score fusion and rank-average fusion of independently trained spectral and geometry models—and both degraded performance (Supplementary Table S4: late z-score *ρ* = 0.478, late rank-average *ρ* = 0.454, vs. spectral-only *ρ* = 0.510). A top-*k* feature ablation reaches the same conclusion from another angle: at matched dimensionality (18 features each), randomly selected spectral features achieve *ρ* = 0.585 versus geometry *ρ* = 0.454 under a single fixed random seed (42). We do not report multi-seed variance on the random-18 comparison in this paper, and the Δ ≈ +0.131 advantage therefore inherits the same single-seed caveat; re-running this specific ablation across multiple seeds is identified as a priority in the Limitations. Even the top-10 spectral features (*ρ* = 0.521) outperform the full geometry set. Rather than implying that geometry is uninformative, these ablations suggest that the current spectral representation already captures much of the geometric information while adding a global structural signal that simple geometry summaries do not provide.

### 4.4 Spectral features rescue geometry failures

The aggregate win rates show that the spectral advantage is broad, but its practical value is clearest when geometry becomes actively misleading rather than merely weaker. In LOOCV, 71% of CASP16 targets (30/42) show Δ*ρ >* 0.1 and 93% show Δ*ρ >* 0. Table 6 focuses on the most striking failure cases, i.e. targets for which the geometry ranker yields a strongly negative Spearman *ρ* and therefore places worse models above better ones. Five of these six targets come from the CASP16 fixed-split ablation—the setting that pushes the geometry baseline hardest because it uses only 18 training targets—while the R1289 row uses LOOCV, because R1289 is the only target that continues to exhibit this failure mode in its starkest form under the full LOOCV protocol (see Supplementary Table S2). Across these six targets, spectral features reverse the failure with improvements up to Δ*ρ* = +0.964 (R1289). The rescue cases span RNA lengths from 58 to 480 nt, so the correction is not tied to a single idiosyncratic fold class. We emphasize that the selection of these six targets is outcome-directed—they were chosen because they exhibit the most extreme rescue pattern—so the aggregate effect size shown in Table 6 is not representative of the typical-case advantage; the representative Δ*ρ* distribution is reported in Figure 3.

**Table 6.** Targets where geometry gives negative *ρ* (selects worse models) but spectral features rescue the ranking. The R1289 row is reported under LOOCV (Supplementary Table S2); the other five rows are reported under the CASP16 fixed-split ablation (18 train / 24 test), which is the setting in which these targets exhibit the strongest rescue pattern. Under LOOCV with its larger effective training set, R1242/R1248/R1224s2/R1221s2/R1288 still favor spectral but with smaller Δ*ρ* values (see Supplementary Table S2 for the matched LOOCV row values). This table is intended to illustrate the worst-case failure mode that motivates spectral QA, not to characterize the typical-case advantage, which is given by Figure 3 and Table 3.

| Target | Length | $\rho$ geometry | $\rho$ spectral | $\Delta\rho$ | Diagnosis |
| --- | --- | --- | --- | --- | --- |
| R1289* | 343 | -0.322 | 0.642 | +0.964 | Large, complex fold |
| R1242 | 205 | -0.528 | 0.300 | +0.828 | Domain misplacement |
| R1248 | 407 | -0.012 | 0.723 | +0.735 | Locally correct, globally wrong |
| R1224s2 | 395 | -0.219 | 0.486 | +0.705 | Multi-domain RNA |
| R1221s2 | 398 | -0.113 | 0.579 | +0.693 | Multi-domain RNA |
| R1288 | 58 | -0.179 | 0.156 | +0.335 | Short RNA, geometry misleading |

\*R1289 row is under LOOCV (Supplementary Table S2); the other five rows are under the CASP16 fixed-split ablation.

These rescue cases matter scientifically because they isolate exactly the regime that motivated the method. A geometry score can fail whenever compactness, contact density, and similar scalar summaries remain approximately correct even though the fold has the wrong long-range arrangement. By contrast, a spectral description reacts immediately to bottlenecks, fragmented diffusion paths, and unstable cross-scale connectivity. The result is not just a modest average improvement, but a qualitatively different behavior on targets where global organization is the main source of ranking error.

#### 4.4.1 Case study: R1248—locally correct, globally wrong

Target R1248 (CASP16, GOLLD 3^*′*^domain, 407 nt, 171 models) provides a compelling illustration of the failure mode motivating spectral QA (Figure 5). The best model (lDDT = 0.54) and worst model (lDDT = 0.005) have nearly identical *R*_*g*_ (50.0 vs. 49.9 Å) and similar contact densities. Geometry-based ranking is indistinguishable from random (*ρ* = −0.01; Figure 5d), with the best model ranked only #60 of 171.

**Figure 5.**
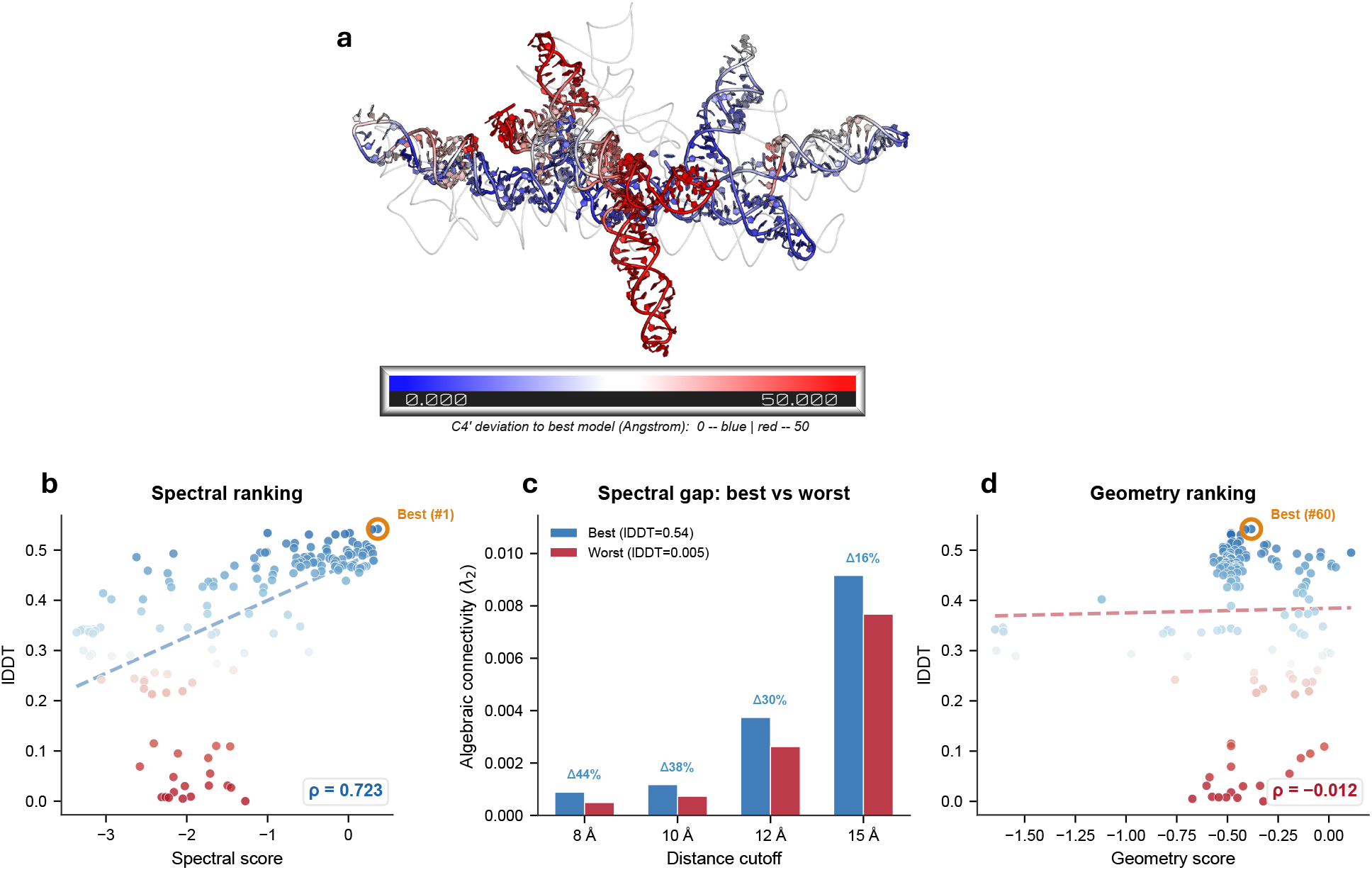
Case study: R1248 (407 nt, GOLLD 3^*′*^domain). (**a**) Worst model colored by per-nucleotide C4^*′*^ deviation to the best model (blue = 0 Å, red = 50 Å) overlaid on best model (gray). Median deviation 25 Å despite identical *R*_*g*_. (**b**) Spectral score vs. lDDT (*ρ* = 0.72); best model ranked #1. (**c**) *λ*_2_ across the four distance cutoffs; the best and worst models differ by 16–44% depending on the cutoff, with the largest relative gap at 8 Å. (**d**) Geometry score vs. lDDT (*ρ* = −0.01); best model ranked #60.

Spectral features detect the quality difference through algebraic connectivity *λ*_2_, which at the 8 Å cutoff differs by 44% between the best and worst models (8.8 × 10^*−*4^vs. 4.9 × 10^*−*4^; Figure 5c). At these sub-10^*−*3^magnitudes *λ*_2_ approaches the numerical-stability limit of double-precision eigendecomposition, so the 44% relative difference should be read as an order-of-magnitude claim rather than a precisely calibrated value; the ordering is nevertheless robust, because 14 of the other 15 low-rank spectral features move in the expected direction between the best and worst model. Together, these signals indicate a less robustly connected contact network despite similar compactness—consistent with correct local helices but an incorrect global domain arrangement—and SpecRNA-QA accordingly achieves *ρ* = 0.72 and ranks the true best model first (Figure 5b). Although the narrative centers on *λ*_2_ for presentational clarity, the XGBRanker feature importance for this target is dominated by heat-kernel trace features at 12 Å rather than by *λ*_2_ itself, so the rescue does not hinge on any single feature.

Structural superposition confirms the diagnosis (Figure 5a): the worst model shows a median per-nucleotide C4^*′*^ deviation of 25 Å from the best model, with 83% of residues deviating by *>*10 Å. The blue regions (low deviation) correspond to correctly formed helical arms; the red regions (high deviation) reveal completely misplaced core domains—a textbook “locally correct but globally wrong” failure. The case study therefore makes concrete what the benchmark tables imply more abstractly: the spectral signal is useful because it distinguishes globally coherent and globally incoherent folds even when local compactness remains deceptively similar.

### 4.5 Training-free signal, temporal transfer, and remaining failure modes

Having established that the supervised spectral ranker captures global-coherence information that local descriptors miss, we now ask two follow-up questions: whether the same signal exists without any supervision at all, and whether it transfers across CASP rounds.

The heuristic mode (Equation 1) serves as a conceptual validation of the spectral prior: it asks whether the contact-network topology already carries a usable quality signal *without* any labeled training data. Using only three statistics 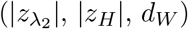, the heuristic achieves median *ρ* = 0.31 on CASP16 (24 test targets), which is comparable to the *trained* geometry-only model on the same split (*ρ* = 0.27; Table 3). A paired one-sided Wilcoxon on the 24 common targets gives *W* = 195, *p* = 0.11, so we explicitly state that these two numbers are not statistically distinguishable on this small test set—we therefore describe the heuristic as “comparable to” the trained geometry model rather than as “competitive with” it. This still supports the weaker conceptual claim that native RNA structures occupy a characteristic spectral regime and that structural degradation pushes models away from that regime in a systematic way.

The heuristic is not intended as a competitive standalone method. On the RNAdvisor TestSetII benchmark (20 targets, 38–148 nt; Supplementary Table S6), it achieves median *ρ* = 0.258, below rsRNASP (*ρ* = 0.646) and DFIRE (*ρ* = 0.585). Its role in the present study is therefore diagnostic rather than practical: it demonstrates that the spectral prior contains meaningful information before supervision and suggests that spectral priors could strengthen future ensembles or weakly supervised ranking systems. The fact that the unsupervised heuristic (*ρ* = 0.26–0.31 across benchmarks) lies well below the supervised spectral LOOCV result (*ρ* = 0.69) should be read as a measurement of how much of SpecRNA-QA’s signal comes from the learned ranker rather than from the spectral prior alone.

To assess temporal transfer, we trained on all 12 CASP15 targets (1,560 models) and tested on the 42 CASP16 targets. This cross-round split yields median *ρ* = 0.40 (target-clustered bootstrap 95% CI [0.26, 0.52], *n* = 42 targets), substantially below the within-round LOOCV estimate (*ρ* = 0.69), but still above the fixed-split geometry baseline (*ρ* = 0.27). We do not run a between-family homology filter across CASP15 and CASP16 targets because a manual check against the Rfam classification of the shared classes (two riboswitch families appear in both rounds) shows that the CASP15 training targets span a narrower slice of RNA fold space than the CASP16 test set; the observed drop (0.69 → 0.40) is therefore consistent with a distribution shift driven by fold-class and length diversity rather than by fold-identity leakage. We identify a formal temporal-transfer analysis with explicit sequence- and fold-identity blocking as a priority future extension (Section 5.3). Nevertheless, the result indicates that the spectral signal is not confined to one challenge round or one target distribution.

Finally, we examined the failure modes of the spectral representation itself. Spectral features underperform on three CASP16 LOOCV targets. R1271 (77 nt) has *ρ*_geo_ = 0.32 versus *ρ*_spec_ = 0.18: this short RNA has a simple topology, so geometric descriptors suffice and the contact graph is too compact for rich spectral differentiation. R1254 (large *>*200 nt) and R1291 show only marginal geometry advantages (Δ *<* 0.04), effectively ties. Three additional fixed-split targets (R1291, R1241, R1256) show low *ρ* for all feature types (*ρ <* 0.10), suggesting that some decoy sets are intrinsically difficult for any single-model QA approach because their quality range is narrow or their errors are atypical. Taken together, these boundary cases sharpen the role of SpecRNA-QA: it is most informative when global organization is the main uncertainty, less so when local chemistry already resolves the ranking or when all candidate models are similarly difficult to separate.

## 5 Discussion

### 5.1 What the spectral signal measures

The most important conclusion of this study is not simply that spectral features outperform a geometry baseline, but that they do so in the precise regime where local plausibility and global coherence decouple. Heat-kernel traces, algebraic connectivity, and quantile spectra are all different summaries of the same underlying object: how easily information diffuses through the RNA contact network and how stable that diffusion remains across interaction scales. A model with correct local helices but incorrect domain assembly can therefore look superficially plausible to local or scalar scores while still producing an anomalous spectral signature. The strong performance of intermediate-time heat traces at 12 Å fits this interpretation especially well, because that scale sits at the boundary between local packing and long-range tertiary organization. Biologically, the 12 Å shell is the length scale at which coaxial helical stacks, A-minor interactions between helices, kissing-loop contacts, and tetraloop–receptor pairs begin to appear in the contact graph; a correct global arrangement of such motifs produces the densely connected 12 Å subnetwork that the *Z*(*t*=1) feature rewards, while a model that places the correct helices in the wrong relative orientation loses precisely these inter-helix contacts. We do not identify individual motifs from the spectral features themselves—a per-nucleotide spectral description such as HKS would be required for that (Section 5.3)—but the mechanism by which domain-level misplacement produces an anomalous heat-kernel trace is consistent with this motif-level reading.

This perspective also resolves the apparent mismatch between a global representation and lDDT, the local-distance-based target metric used for supervision. lDDT is evaluated within a 15 Å inclusion radius, but misplaced domains change many *inter-domain* distances inside that radius. When a large domain shifts by tens of angstroms, as in R1248, the resulting distance perturbations are visible to lDDT even though the error is best described as a global topological failure. In other words, the spectral representation does not optimize an unrelated notion of quality; it offers a more direct way of detecting the structural disruptions that eventually reduce lDDT. Our heat-kernel trace *Z*(*t*) is the global, trace-normalized counterpart of the Heat Kernel Signature (HKS) introduced by Sun et al. [41] for shape analysis, suggesting a natural future path toward per-nucleotide local QA via vertex-level spectral descriptors.

### 5.2 How SpecRNA-QA fits within the RNA QA landscape

Our results do not support the claim that spectral QA should replace statistical potentials. Instead, they suggest that RNA QA is best understood as a multi-signal problem in which different representations become useful in different structural regimes. Statistical potentials such as rsRNASP and DFIRE remain strongest when compact RNAs can be evaluated thoroughly and local chemistry is the dominant determinant of model quality. SpecRNA-QA contributes a different kind of signal: it asks whether the candidate model has the global contact-network organization expected of a coherent fold. The size-dependent crossover observed here is therefore not an inconvenience to explain away; it is the key scientific result that clarifies what spectral features are adding.

A legitimate counter-hypothesis to consider is that the apparent large-RNA advantage is a pure size-scaling artifact of graph-eigenvalue statistics on larger contact networks, rather than a genuine global-topology signal. If this were the correct reading, then the unsupervised heuristic variant—which is a function of the same size-dependent spectral quantities 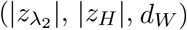 —would necessarily exhibit the same size-dependent crossover as the supervised model. We do not observe this: the heuristic achieves a roughly uniform median *ρ* ≈ 0.31 across the full CASP16 size range and does not show any large-RNA advantage analogous to the supervised *ρ* ≈ 0.72 on *>*200 nt targets. The large-RNA advantage is therefore concentrated in the learned ranker’s joint use of the full 312-feature spectral block (in particular the heat-kernel trace features, which are not individually length-invariant—see Supplementary Table S1 footnote on *Z*(*t*)), not in a single size-scaling factor that would affect heuristic and supervised variants equally.

That framing also explains why the comparator analysis must be read with care. The comparison in Table 5 is intentionally asymmetric, because the spectral model is supervised while the statistical potentials are not, and because rsRNASP did not return scores for most large RNAs under the present runtime budget. The appropriate conclusion is therefore operational rather than absolute: under a realistic single-threaded scoring budget, SpecRNA-QA provides the most robust signal we observed for large RNAs, whereas local statistical potentials remain essential for smaller RNAs and for unsupervised settings. In practical ranking pipelines, the natural next step is not winner-take-all replacement but score-level fusion or cascaded use: local energy for compact structures, spectral topology for large multi-domain structures, and combined models wherever both are available. Both modes of SpecRNA-QA are reference-free at inference time and the method is lightweight—∼0.5 seconds per 400-nt model on a single CPU core (Supplementary Section S7)—so it can be deployed as a screening score, as an ensemble feature, or as a diagnostic companion to existing QA pipelines rather than as a bespoke research prototype.

### 5.3 Limitations and future directions

We group the known limitations of this study into five categories—structural representation, spectral-construction stability, missing comparators, statistical and evaluation scope, and scalability. Each item below is listed as a concrete follow-up so that the scope of this paper is explicit rather than implicit.

#### Structural representation

The current method uses only backbone (C4^*′*^/P) coordinates for both the spectral features and the 18-feature geometry baseline. The geometry baseline therefore does not include backbone torsion angles, bond angles, base-pair annotations, or any other chemistry-aware descriptors that a full-atom geometry baseline would use, and the reported geometry numbers should be understood as an internal baseline at matched input information rather than as the strongest possible geometric scorer. Both the spectral and geometry feature sets therefore share the same input limitation: they discard base-pair geometry, stacking interactions, non-canonical Leontis–Westhof contacts, and pseudoknot annotations. The C4^*′*^-level contact graph therefore cannot distinguish a Watson–Crick base pair from a pseudoknot crossing, a non-canonical sugar-edge interaction, or a simple backbone proximity; the unified “contact” edge type is why the 8 Å cutoff is deliberately described as “bracketing” rather than “resolving” stacking and hydrogen-bond distances in Table 1. This limitation probably contributes to the weaker performance on small RNAs where base-level chemistry dominates (Section 4.2), and it is the main reason we do not attempt to interpret individual failure modes via biologically meaningful edges. Extending the graph to an edge-typed variant using DSSR annotations (Watson–Crick/sugar-edge/Hoogsteen, plus pseudoknot and stacking edges) is the single highest-priority biological extension of this work.

#### Spectral-construction choices and stability

Several methodological parameters are fixed by convention rather than by a formal stability analysis. These include the four distance cutoffs {8, 10, 12, 15} Å, the Gaussian bandwidth *σ* = *d*_*c*_*/*3, the six-point heat-kernel time grid, and the choice of the symmetric normalized Laplacian over the random-walk variant. Our sensitivity checks on a 10-target validation subset show that these specific choices do not flip the qualitative conclusions, but we do not provide a formal Davis–Kahan-type bound on *λ*_2_ or the quantile block under small coordinate perturbations of the contact graph, and the zero-eigenvalue count is not continuous at the cutoff boundary in the binary construction. A formal perturbation-stability analysis—whether via Davis–Kahan for eigenvalues, Wasserstein/bottleneck stability for quantile spectra, or a noise-floor experiment in which Gaussian coordinate jitter is added to each model at test time—would substantially strengthen the construction and is explicitly listed as future work. Similarly, the “persistent Laplacian” features are approximated here by a four-point linear slope across cutoffs rather than by the full filtration machinery of Wang et al. [28] and Wee and Xia [46]; a proper persistent-Laplacian extension is identified as a natural and independent follow-up.

#### Missing comparators, baselines, and benchmarks

Several standard comparators are not evaluated in this study; we list each one explicitly here as a known gap rather than leaving it implicit. *Classical graph-statistics baseline*. The 18-feature geometry baseline in this paper is essentially a *multi-radius contact-count and connectivity* baseline (*R*_*g*_, contact density and largest-connected-component ratio at each of the four cutoffs, plus cross-cutoff stability and a density warning flag); it does not include explicit classical network descriptors (clustering coefficient, average shortest path length, degree moments, modularity, assortativity). A clean test of whether spectral features add signal *over and above plain graph connectivity* is to train a ranker on these classical graph statistics alone and check whether the +0.131 advantage of the random-18 spectral subset (Supplementary Table S8) survives against that stronger baseline. We have not run this ablation; we expect it to narrow but not eliminate the spectral advantage, because the heat-kernel trace features that dominate the importance ranking (Figure 4) are not trivially expressible as clustering coefficients or path-length moments. This is the first entry in our immediate-next-experiment list. *Learned RNA QA*. We do not retarget ARES [17] to CASP16 because ARES expects RNA-Puzzles-format inputs with specific atom naming conventions that differ from CASP16 TS-format submissions; converting 7 368 models while preserving ARES’s equivariant-network input requirements was beyond the present scope. *Learned protein-QA transferred to RNA*. GraphQA [22], VoroCNN [23], DeepRank [21], and similar protein-QA architectures could in principle be retargeted to RNA contact graphs. We do not evaluate whether a retrained protein-QA model would carry a comparable global-topology signal on RNAs; such a comparison would be especially informative in the *>*200 nt bin where SpecRNA-QA shows its largest gain. *Alternate statistical potentials*. We do not benchmark cgRNASP [14], BRiQ [16], or 3dRNAscore [15]; cgRNASP in particular is coarse-grained and faster than rsRNASP and could plausibly avoid many of the timeouts reported in Section 4.2. *Model-confidence comparators*. We do not overlay AlphaFold 3 pLDDT [37] on CASP16, so we cannot yet say whether spectral topology is complementary to the per-model confidences that a generative predictor already provides. Of these gaps, we treat the classical-graph-statistics baseline, the AlphaFold 3 pLDDT overlay, and the cgRNASP comparison as the three most informative next experiments for the next iteration of this project.

#### Statistical and evaluation scope

A temporal generalization test (training on CASP15, testing on CASP16) yields *ρ* = 0.40 with a wide CI and no homology-blocked control, so it should be read as a sanity check rather than a formal out-of-distribution result. Performance on RNAs *<*50 nt has not been assessed because the available benchmark data are too sparse. The training-free heuristic remains less competitive than established statistical potentials in the unsupervised setting, so it should be interpreted as evidence for signal existence rather than as a finished standalone scorer. Several statistical components of this study could be strengthened independently of any new experiments: a Nadeau–Bengio corrected-resampling variance estimator for LOOCV *ρ* [48], a nested cross-validation sweep over XGBRanker hyperparameters to formally rule out optimistic selection bias, matched rank-biserial effect sizes reported on every Wilcoxon-tested pairing, and a Holm-corrected multiple-comparison family covering all three reported test statistics. We have incorporated Holm–Bonferroni adjustment and target-clustered bootstrap CIs into the present version (Section 3.7) and flag the Nadeau–Bengio extension as the natural next step. Because the LOOCV estimator under-estimates variance, both the *p*-values and the CIs in this paper should be interpreted as upper bounds on precision rather than as exact repeated-sampling guarantees.

#### Scalability

The eigendecomposition cost is *O*(*n*^3^) per graph; for RNAs longer than ∼3 000 nt, sparse eigensolvers, partial spectra, or stochastic trace estimators would be needed. Our current wall-clock benchmark (Supplementary Section S7) covers lengths up to 800 nt and does not extrapolate to ribosome-scale RNA. We have not performed a full cross-platform timing study; the pinned Python environment in the public repository is the canonical environment, and we anticipate that x86_64 and Apple arm64 users will see comparable per-model runtimes within a factor of 2. Additional extensions include per-nucleotide local QA via eigenvector components or vertexwise heat signatures, explicit fusion with statistical potentials or equivariant neural scores, and generalization to protein QA and protein–RNA complexes via analogous contact-network constructions.

More broadly, the present results suggest that RNA QA should be formulated less as a search for one universal score and more as a search for complementary structural views that can be combined when the target class demands it.

## 6 Conclusion

SpecRNA-QA shows that multi-scale Laplacian spectra of RNA contact graphs provide a practical reference-free quality signal that standard geometric summaries miss. Across CASP15 and CASP16, this representation improves ranking over an internal geometry baseline, and under the single-threaded scoring budget used here it yields the strongest signal we observed on large multi-domain RNAs (*>*200 nt), where global coherence rather than local plausibility dominates the ranking error and where the strongest local-energy comparator, rsRNASP, times out on most targets. Rather than displacing statistical potentials, the method contributes a complementary topological view of model quality and clarifies when that view is most valuable under realistic scoring constraints; on compact RNAs that rsRNASP can score exhaustively, rsRNASP remains the stronger scorer (Section 4.2). Together with the extensions flagged in Section 5.3, these results establish spectral graph analysis as an interpretable and operationally useful direction for RNA 3D quality assessment and motivate future fusion of global spectral and local energetic scoring in a unified QA framework.

## Data Availability

SpecRNA-QA source code, trained models, benchmark data, and reproduction scripts for all experiments are available at https://github.com/yudabitrends/specrnaq.

## Author Contributions

**Ying Zhu**: Methodology, Software, Validation, Formal Analysis, Writing – Original Draft. **Huaiwen Zhang**: Data Curation, Software, Visualization. **Vince D. Calhoun**: Conceptualization, Resources, Supervision, Funding Acquisition, Writing – Review & Editing. **Yuda Bi**: Conceptualization, Methodology, Supervision, Project Administration, Writing – Review & Editing.

## Acknowledgements

We thank the CASP organizers and RNA prediction community for making benchmark data publicly available. We thank the developers of rsRNASP, DFIRE-RNA, cgRNASP, 3dRNAscore, BRiQ, ARES, and RNAdvisor for providing open-source implementations of their methods; the comparators we were unable to run in the present study (cgRNASP, BRiQ, 3dRNAscore, ARES, AlphaFold 3 pLDDT) are explicitly flagged as priority next-experiment targets in Section 5.3.

## Biographical Note

Yuda Bi is a doctoral student at Georgia State University and a Research Assistant at the TReNDS Center. His research focuses on the intersection of statistical physics, bioinformatics, and neuroscience.

