## supplementary for "Spectral Graph Features for Reference-free RNA 3D Quality Assessment"

### Supplementary Materials

Spectral Graph Features Capture Global Topology for
Reference-free RNA 3D Structure Quality Assessment

#### S1 Spectral Feature Catalog

Table **??** provides the complete catalog of features extracted by SpecRNA-QA. For each of the 8 contact graphs (4 distance cutoffs $\times$ 2 kernel types), the spectral feature block produces 33 features. Combined with cross-scale stability statistics, native prior features, and geometric baseline features, the total is 332 features per structure (314 spectral + 18 geometric).

Table S1: Complete spectral feature catalog. Each feature block is computed per graph (4 cutoffs × 2 kernels = 8 graphs). Cross-scale statistics are computed across the 4 cutoffs for each kernel type.

| Category | Features | Count/graph | Physical meaning |
|---|---|---|---|
| Low eigenvalues | $\lambda_2$–$\lambda_6$ | 5 | Algebraic connectivity and dominant structural modes |
| Quantile spectrum | $q_{0.05}$–$q_{0.95}$ (11 percentiles) | 11 | Eigenvalue distribution shape, length-invariant |
| Spectral entropy | $H = -\sum \tilde{\lambda}_i \log \tilde{\lambda}_i$ | 1 | Complexity of spectral distribution |
| Spectral moments | $\mu_1$–$\mu_4$ | 4 | Mean, variance, skewness, kurtosis |
| Heat-kernel trace | $Z(t)$ at $t \in \{0.1, 0.5, 1, 2, 5, 10\}$ | 6 | Multi-scale diffusion probe (local→global) |
| Inverse participation ratio | IPR of 5 lowest modes | 5 | Mode localization |
| Zero eigenvalue count | $n_0$ | 1 | Number of connected components |
| **Per-graph subtotal** | | **33** | |
| **× 8 graphs** | | **264** | |
| Cross-scale stability | Mean, var., slope across 4 cutoffs: binary $\lambda_2$ + $H$ (6); Gaussian $\lambda_2$ + $H$ + 11 quantiles (39) | 45 | Robustness across distance scales |
| Native prior features | $d_W$, $z_{\lambda_2}$, $z_H$ | 3 | Distance to native spectral template |
| **Spectral total** | | **314** | |
| Geometric baseline | Length, $R_g$, missing fraction | 3 | Basic structural descriptors |
| | Contact density (×4 cutoffs) | 4 | Local packing density |
| | Largest component ratio (×4) | 4 | Graph connectivity |
| | Cross-cutoff stability (×6) | 6 | |
| | Density warning flag | 1 | |
| **Geometric total** | | **18** | |
| **Grand total** | | **332** | |

#### S2 Per-target LOOCV Results

Tables **??** and **??** report per-target Spearman $\rho$ for all targets under leave-one-out cross-validation, comparing spectral-only and geometry-only feature sets.

Table S2: Per-target LOOCV results on CASP16 (42 targets). $\Delta\rho = \rho_{\text{spectral}} - \rho_{\text{geometry}}$. Targets sorted by $\Delta\rho$ descending.

| Target | Size bin | Models | $\rho$ spectral | $\rho$ geometry | $\Delta\rho$ | Winner |
|---|---|---|---|---|---|---|
| R1289 | >200 | 157 | 0.642 | −0.322 | +0.964 | spectral |
| R1248 | >200 | 171 | 0.757 | 0.280 | +0.477 | spectral |

| Target | Size bin | Models | $\rho$ spectral | $\rho$ geometry | $\Delta\rho$ | Winner |
|---|---|---|---|---|---|---|
| R1252 | >200 | 125 | 0.682 | 0.106 | +0.576 | spectral |
| R1212 | >200 | 201 | 0.491 | 0.089 | +0.402 | spectral |
| R1281 | >200 | 164 | 0.505 | 0.449 | +0.056 | spectral |
| R1224s2 | >200 | 197 | 0.763 | 0.242 | +0.521 | spectral |
| R1221s2 | >200 | 189 | 0.711 | 0.529 | +0.182 | spectral |
| R1283 | >200 | 156 | 0.770 | 0.609 | +0.161 | spectral |
| R1285 | >200 | 155 | 0.754 | 0.452 | +0.302 | spectral |
| R1286 | >200 | 162 | 0.809 | 0.663 | +0.146 | spectral |
| R0250 | >200 | 164 | 0.748 | 0.538 | +0.210 | spectral |
| R0251 | >200 | 161 | 0.759 | 0.503 | +0.256 | spectral |
| R0252 | >200 | 162 | 0.808 | 0.527 | +0.281 | spectral |
| R0253 | >200 | 156 | 0.781 | 0.598 | +0.183 | spectral |
| R0254 | >200 | 162 | 0.751 | 0.524 | +0.228 | spectral |
| R0281 | >200 | 155 | 0.730 | 0.384 | +0.346 | spectral |
| R0283 | >200 | 155 | 0.770 | 0.468 | +0.301 | spectral |
| R0285 | >200 | 171 | 0.723 | 0.677 | +0.046 | spectral |
| R0290 | >200 | 140 | 0.522 | 0.422 | +0.099 | spectral |
| R1241 | >200 | 179 | 0.688 | 0.462 | +0.226 | spectral |
| R1242 | >200 | 191 | 0.520 | 0.343 | +0.177 | spectral |
| R1250 | >200 | 146 | 0.714 | 0.377 | +0.336 | spectral |
| R1251 | >200 | 129 | 0.695 | 0.599 | +0.096 | spectral |
| R1254 | >200 | 83 | 0.675 | 0.708 | −0.032 | geometry |
| R1290 | >200 | 165 | 0.680 | 0.567 | +0.113 | spectral |
| R1291 | >200 | 174 | 0.641 | 0.645 | −0.003 | geometry |
| R1203 | 100–200 | 183 | 0.689 | 0.302 | +0.388 | spectral |
| R1255 | 100–200 | 189 | 0.486 | 0.202 | +0.283 | spectral |
| R1256 | 100–200 | 187 | 0.371 | 0.282 | +0.090 | spectral |
| R1205 | 50–100 | 186 | 0.276 | −0.120 | +0.397 | spectral |
| R1209 | 50–100 | 212 | 0.485 | 0.468 | +0.017 | spectral |
| R1211 | 50–100 | 184 | 0.391 | −0.006 | +0.398 | spectral |
| R1221s3 | 50–100 | 184 | 0.652 | 0.504 | +0.148 | spectral |
| R1224s3 | 50–100 | 198 | 0.757 | 0.668 | +0.090 | spectral |
| R1261 | 50–100 | 227 | 0.731 | 0.664 | +0.067 | spectral |
| R1262 | 50–100 | 216 | 0.757 | 0.622 | +0.135 | spectral |
| R1263 | 50–100 | 217 | 0.549 | 0.444 | +0.105 | spectral |
| R1264 | 50–100 | 206 | 0.646 | 0.616 | +0.030 | spectral |
| R1271 | 50–100 | 172 | 0.184 | 0.323 | −0.139 | geometry |
| R1288 | 50–100 | 193 | 0.107 | −0.027 | +0.135 | spectral |
| R1293 | 50–100 | 214 | 0.591 | 0.403 | +0.187 | spectral |
| R1296 | 50–100 | 230 | 0.744 | 0.634 | +0.110 | spectral |

Table S3: Per-target LOOCV results on CASP15 (12 targets).

| **Target** | **Size bin** | **Models** | **$\rho$ spectral** | **$\rho$ geometry** | **$\Delta\rho$** | **Winner** |
|---|---|---|---|---|---|---|
| R1107 | 50–100 | 131 | 0.787 | 0.762 | +0.025 | spectral |
| R1108 | 50–100 | 115 | 0.812 | 0.613 | +0.198 | spectral |
| R1116 | 100–200 | 145 | 0.630 | 0.445 | +0.186 | spectral |
| R1117 | <50 | 54 | 0.500 | −0.057 | +0.557 | spectral |
| R1126 | >200 | 140 | 0.224 | 0.102 | +0.122 | spectral |
| R1128 | >200 | 137 | 0.349 | −0.075 | +0.425 | spectral |
| R1136 | >200 | 158 | 0.802 | 0.505 | +0.296 | spectral |
| R1138 | >200 | 130 | 0.708 | −0.295 | +1.003 | spectral |
| R1149 | 100–200 | 138 | 0.627 | 0.494 | +0.133 | spectral |
| R1156 | 100–200 | 145 | 0.625 | 0.477 | +0.148 | spectral |
| R1189 | 100–200 | 136 | 0.710 | 0.348 | +0.362 | spectral |
| R1190 | 100–200 | 131 | 0.628 | 0.309 | +0.319 | spectral |

### S3 Late Fusion Comparison

Table **??** compares feature combination strategies on the CASP16 fixed-split benchmark (24 test targets).

Table S4: Feature combination strategies on CASP16 (24 test targets, fixed split).

| **Strategy** | **Features used** | **Median $\rho$** | **Top-1 hit** |
|---|---|---|---|
| Spectral only (XGBRanker) | 314 spectral | **0.510** | **20.8%** |
| Early concatenation (all) | 332 all | 0.497 | 20.8% |
| Late fusion (z-score) | spectral + geometry scores | 0.478 | 4.2% |
| Late fusion (rank average) | spectral + geometry ranks | 0.454 | 4.2% |
| Geometry only (XGBRanker) | 18 geometric | 0.265 | 4.2% |

### S4 XGBRanker Hyperparameters

Table **??** lists the hyperparameters used for the XGBRanker model in all experiments.

Table S5: XGBRanker hyperparameters.

| **Parameter** | **Value** | **Rationale** |
|---|---|---|
| Objective | rank:pairwise | LambdaMART pairwise ranking loss |
| n_estimators | 240 | Sufficient convergence without overfitting |
| learning_rate | 0.05 | Conservative step size |
| max_depth | 4 | Moderate interaction complexity |
| min_child_weight | 1.0 | Default; allows fine splits |
| subsample | 0.9 | Per-iteration row sampling |
| colsample_bytree | 0.8 | Feature subsampling per tree |
| reg_lambda | 1.0 | L2 regularization |
| random_state | 42 | Reproducibility |

#### S5 RNAdvisor TestSetII Detailed Results

Table ?? reports per-target Spearman $\rho$ on the RNAdvisor TestSetII benchmark (20 targets, 38–148 nt, ∼40 decoys per target). SpecRNA-QA operates in training-free heuristic mode using only three spectral statistics.

Table S6: Per-target comparison on RNAdvisor TestSetII. Bold indicates best among reference-free methods (excluding TM-SCORE which requires a native structure).

| **Target** | **Length** | **SpecRNA-QA** | **rsRNASP** | **DFIRE** | **RASP** |
|---|---|---|---|---|---|
| 1Z43 | 101 | 0.284 | **0.596** | 0.496 | 0.297 |
| 3A3A | 86 | 0.359 | **0.812** | 0.693 | 0.579 |
| 3IVN | 69 | 0.243 | **0.513** | 0.530 | 0.135 |
| 3L0U | 73 | −0.271 | **0.737** | 0.670 | −0.402 |
| 3LA5 | 71 | 0.422 | **0.886** | 0.837 | −0.317 |
| 3RKF | 67 | 0.385 | **0.931** | 0.928 | 0.498 |
| 3SKI | 68 | −0.069 | **0.894** | 0.769 | 0.281 |
| 4AOB | 94 | **0.746** | 0.860 | 0.758 | −0.187 |
| 4FEN | 67 | 0.097 | **0.924** | 0.854 | −0.180 |
| 5D5L | 77 | 0.344 | **0.719** | 0.592 | 0.078 |
| 5FJC | 93 | **0.757** | 0.903 | 0.897 | 0.067 |
| 5SWD | 65 | −0.001 | **0.477** | 0.149 | 0.244 |
| 6C27 | 47 | −0.023 | **0.261** | 0.305 | 0.327 |
| 6E8S | 38 | 0.274 | **0.471** | 0.474 | 0.152 |
| 6JQ6 | 81 | **0.347** | 0.325 | 0.236 | 0.295 |
| 6TFE | 52 | **0.662** | 0.506 | 0.591 | −0.309 |
| 6VMY | 148 | −0.087 | **0.531** | 0.341 | 0.244 |
| 7D82 | 50 | 0.042 | **0.623** | 0.495 | 0.199 |
| 7K16 | 51 | 0.157 | **0.669** | 0.579 | 0.247 |
| 7KJU | 75 | 0.126 | 0.334 | 0.071 | **0.352** |
| **Median** | | **0.258** | **0.646** | **0.585** | **0.221** |

#### S6 Pooled vs. Median Spearman Correlation

The *pooled Spearman* $\rho$ computes a single rank correlation across all models from all targets combined. For RNA QA benchmarks, this metric is problematic because:

1. **Score incomparability**: Predicted QA scores are meaningful only within a single target. A model with score 0.5 for a 400-nt target and one with score 0.5 for a 60-nt target are not comparable—they correspond to different quality levels.
2. **Length bias**: Targets vary from 29 to 833 nt. Longer targets with more models (up to 230) dominate the pooled metric, while short targets with fewer models contribute little.
3. **Distribution heterogeneity**: lDDT distributions differ markedly across targets. Some have clustered quality (narrow distribution), others have wide spreads. Pooling conflates these fundamentally different ranking tasks.

The *median per-target Spearman* $\rho$ avoids these issues by computing $\rho$ independently within each target, then taking the median. This reflects the question "how well does the method rank models for a typical target?" and gives equal weight to easy and hard targets regardless of model count. We follow the recommendation of Kretsch et al. (2025) in adopting this as the primary metric.

For completeness, we report both metrics throughout the paper and supplementary tables. On CASP16 LOOCV, spectral-only achieves pooled $\rho = 0.432$ vs. geometry 0.238—a consistent advantage, though the absolute values are diluted by cross-target pooling.

### S7 Computational Cost

SpecRNA-QA's computational cost is dominated by eigendecomposition of $n \times n$ Laplacian matrices, where $n$ is the number of nucleotides. For each structure, 8 eigendecompositions are performed (4 distance cutoffs $\times$ 2 kernel types).

Table S7: Representative runtime on Apple Silicon (M-series, single CPU core).

| **Operation** | **100 nt** | **400 nt** | **800 nt** |
|---|---|---|---|
| Distance matrix | $<$1 ms | $\sim$5 ms | $\sim$20 ms |
| 8 graph constructions | $\sim$2 ms | $\sim$30 ms | $\sim$120 ms |
| 8 eigendecompositions | $\sim$10 ms | $\sim$500 ms | $\sim$4 s |
| Feature extraction | $\sim$1 ms | $\sim$5 ms | $\sim$10 ms |
| Prior enrichment | $<$1 ms | $<$1 ms | $<$1 ms |
| XGBRanker prediction | $<$1 ms | $<$1 ms | $<$1 ms |
| **Total per model** | **$\sim$15 ms** | **$\sim$0.5 s** | **$\sim$4.2 s** |

The full CASP16 benchmark (7,368 models, 42 targets) completes in $\sim$25 minutes with feature caching across LOOCV folds. Memory usage is $O(n^2)$ per graph, peaking at $\sim$5 MB for 800-nt RNA. No GPU is required.

### S8 Top-$k$ Feature Ablation

To directly control for the dimensionality confound (314 spectral vs. 18 geometric features), we trained XGBRanker in LOOCV using only the top-$k$ spectral features ranked by importance (Table **??**). Additionally, we compared 18 randomly selected spectral features against the 18 geometry features at matched dimensionality.

Table S8: Top-$k$ feature ablation on CASP16 LOOCV (42 targets). Features ranked by XGBRanker gain-based importance. Random-18-spectral uses a fixed random seed (42) for reproducibility. Note: these ablation results use XGBRanker directly without the full SpecRNAQAModel wrapper (which includes z-score normalization and isotonic calibration), so absolute $\rho$ values are slightly lower than in Table 1 of the main text (e.g., all-spectral $\rho = 0.661$ here vs. 0.689 in LOOCV). Relative comparisons between configurations are valid.

| **Configuration** | **Features** | **Median $\rho$** |
|---|---|---|
| Geometry (baseline) | 18 | 0.454 |
| Top-10 spectral | 10 | 0.521 |
| Random 18 spectral | 18 | 0.585 |
| Top-20 spectral | 20 | 0.620 |
| Top-30 spectral | 30 | 0.647 |
| Top-50 spectral | 50 | 0.647 |
| Top-100 spectral | 100 | 0.652 |
| All spectral | 314 | 0.661 |
| All features (spectral + geometry) | 332 | 0.681 |

At matched dimensionality (18 features), randomly selected spectral features ($\rho = 0.585$) substantially outperform the geometry baseline ($\rho = 0.454$), a $\Delta = +0.131$ advantage that cannot be explained by feature count alone. Even the top-10 spectral features ($\rho = 0.521$) outperform the 18-feature geometry set with fewer dimensions. Performance plateaus around $k = 30$–$50$, suggesting that a compact set of spectral descriptors—dominated by heat-kernel traces and quantile spectrum features—captures most of the quality-discriminating information.
